# Mechanistic insights into NAD synthase NMNAT chaperoning phosphorylated Tau from pathological aggregation

**DOI:** 10.1101/800565

**Authors:** Xiaojuan Ma, Jinxia Lu, Yi Zhu, Jingfei Xie, Chong Li, Austin Shin, Jiali Qiang, Jiaqi Liu, Shuai Dou, Yi Xiao, Chuchu Wang, Chunyu Jia, Houfang Long, Yanshan Fang, Lin Jiang, Yaoyang Zhang, Shengnan Zhang, R. Grace Zhai, Cong Liu, Dan Li

## Abstract

Tau hyper-phosphorylation and deposition into neurofibrillary tangles have been found in brains of patients with Alzheimer’s disease (AD) and other tauopathies. Molecular chaperones are involved in regulating the pathological aggregation of phosphorylated Tau (pTau) and modulating disease progression. Here, we report that nicotinamide mononucleotide adenylyltransferase (NMNAT), a well-known NAD synthase, serves as a chaperone of pTau to prevent its amyloid aggregation *in vitro* as well as mitigate its pathology in a fly tauopathy model. By combining NMR spectroscopy, crystallography, single-molecule and computational approaches, we revealed that NMNAT adopts its enzymatic pocket to specifically bind the phosphorylated sites of pTau, which can be competitively disrupted by the enzymatic substrates of NMNAT. Moreover, we found that NMNAT serves as a co-chaperone of Hsp90 for the specific recognition of pTau over Tau. Our work uncovers a dedicated chaperone of pTau and suggests NMNAT as a key node between NAD metabolism and Tau homeostasis in aging and neurodegeneration.

## Introduction

Phosphorylated Tau (pTau) is the major component of the neurofibrillary tangles that are commonly found in the brains of patients with Alzheimer’s disease (AD) and many other tauopathy-related neurodegenerative diseases (Avila, 2006; Diane P. Hanger, Anderton, & Noble, 2009; Diane P. Hanger, Betts, Loviny, Blackstock, & Anderton, 2002; D. P. Hanger et al., 1991). Tau is an intrinsically disordered protein with a high abundance in neurons (Hirokawa, Funakoshi, Satoharada, & Kanai, 1996; Konzack, Thies, Marx, Mandelkow, & Mandelkow, 2007). There are six isoforms of Tau in human central nervous system due to alternative splicing (Goedert, Spillantini, Jakes, Rutherford, & Crowther, 1989). Under the physiological condition, Tau associates with microtubules and modulates the stability of axonal microtubules (Drechsel, Hyman, Cobb, & Kirschner, 1992). Whereas, phosphorylation of Tau by protein kinases such as microtubule affinity regulating kinase 2 (MARK2), causes the release of Tau from microtubule binding and proceeds hyper-phosphorylation and aggregation of Tau (Ando et al., 2016; Biernat, Gustke, Drewes, Mandelkow, & Mandelkow, 1993; Drewes, 2004; Drewes, Ebneth, Preuss, Mandelkow, & Mandelkow, 1997; Gu et al., 2013), which is closely associated with the pathogenesis of AD and other tauopathies (Drewes et al., 1997). Different proteins including chaperones (Hsp90, Hsc70/Hsp70) (Dickey, Kamal, et al., 2007), proteasome (Dickey, Patterson, Dickson, & Petrucelli, 2007) and protein phosphatase 2A (PP2A)(Gong, Singh, Grundke-Iqbal, & Iqbal, 1993) were found to play important roles in maintaining Tau homeostasis including preventing abnormal hyper-phosphorylation and aggregation, and facilitating pTau degradation (Diane P. Hanger et al., 2009).

Nicotinamide mononucleotide adenylyltransferase (NMNAT) was initially identified as an NAD synthase that catalyzes the reversible conversion of NMN (nicotinamide mononucleotide) to NAD in the final step of both the *de novo* biosynthesis and salvage pathways in most organisms across all three kingdoms of life (Magni, Amici, Emanuelli, Raffaelli, & Ruggieri, 1999). NMNAT is indispensable in maintaining neuronal homeostasis (Araki, Sasaki, & Milbrandt, 2004; Conforti et al., 2009). Familial mutations of NMNAT have been found to cause Leber congenital amaurosis 9 (LCA9) (Chiang et al., 2012; Falk et al., 2012; Koenekoop et al., 2012; Perrault et al., 2012) and retinal degeneration (Beirowski, Babetto, Coleman, & Martin, 2008). Moreover, NMNAT is closely related to AD and other Tauopathgies. The mRNA level of human NMNAT2 (hN2), one of the three isoforms of human NMNATs (Raffaelli et al., 2002), decreases in patients of AD (Ali et al., 2016). Abundant hN2 proteins were detected in the insoluble fraction of AD patients, which also contains pTau and HSP90 (Ali et al., 2016). In addition, NMNAT plays a protective role in different cellular and animal models of AD (Ali, Li-Kroeger, Bellen, Zhai, & Lu, 2013; Conforti, Gilley, & Coleman, 2014; Fang, Soares, Teng, Geary, & Bonini, 2012; Ocampo, Liu, & Barrientos, 2013). Over-expression of different isoforms of NMNAT can significantly reduce the abnormal aggregation (Zhai et al., 2008) and cytotoxicity of pTau and relieve pTau burden in different models of AD (Rossi et al., 2018) and frontotemporal dementia with parkinsonism linked to chromosome 17 (FTDP-17) (Ali, Ruan, & Zhai, 2012; Ljungberg et al., 2012).

Intriguingly, in addition to its well-studied NAD synthase activity, NMNAT has been found to be able to retrieve the activity of luciferase from heat-denatured amorphous aggregation suggesting a chaperone activity of NMNAT (Ali et al., 2016; Zhai et al., 2008). However, it remains puzzling how a single domain enzyme, at the same time, fulfills a chaperone activity. Moreover, since NAD is an essential cofactor in numerous cellular processes (e.g. transcriptional regulation (D’Amours, Germain, Orth, Dixit, & Poirier, 1998; Shogren-Knaak et al., 2006; T. Zhang et al., 2009) and oxidative reactions (Lewis et al., 2014)), it is confusing whether the protective role of NMNAT in AD animal model comes directly from its potential chaperone activity or indirectly from its enzymatic activity. In this work, we investigated the chaperone activity of NMNAT against the amyloid aggregation of Tau *in vitro* and in the fly model. By combining multiple biophysical and computational approaches, we revealed the molecular mechanism of NMNAT as a specific chaperone of pTau. Our work provides the structural basis for how NMNAT manages its dual functions as both an enzyme and a chaperone, as well as how NMNAT specifically recognizes pTau and serves as a cochaperone of Hsp90 for pTau clearance. Our work suggests an interplay of NAD metabolism and the progression of Tau pathology in aging and neurodegeneration.

## Results

### NMNAT family members exhibit a conserved chaperone activity in preventing pTau aggregation

In the preparation of pTau proteins, we used MARK2 to phosphorylate Tau23 and a truncated construct—K19 **(Figure 1A)** (Gu et al., 2013). he MARK2 phosphorylation sites on Tau23 and K19 were characterized by using 2D ^1^H-^15^N NMR HSQC spectra. Consistent with previous reports (Schwalbe et al., 2013; Timm et al., 2003a), S S262 of R1, S324 of R3, S352 and S356 of R4 were phosphorylated in both MARK2-treated Tau23 and K19. Besides, pTau23 exhibited two additional phosphorylated sites: S413 and S416 **(Figure1A - figure supplement 1)**.

**Figure 1.**
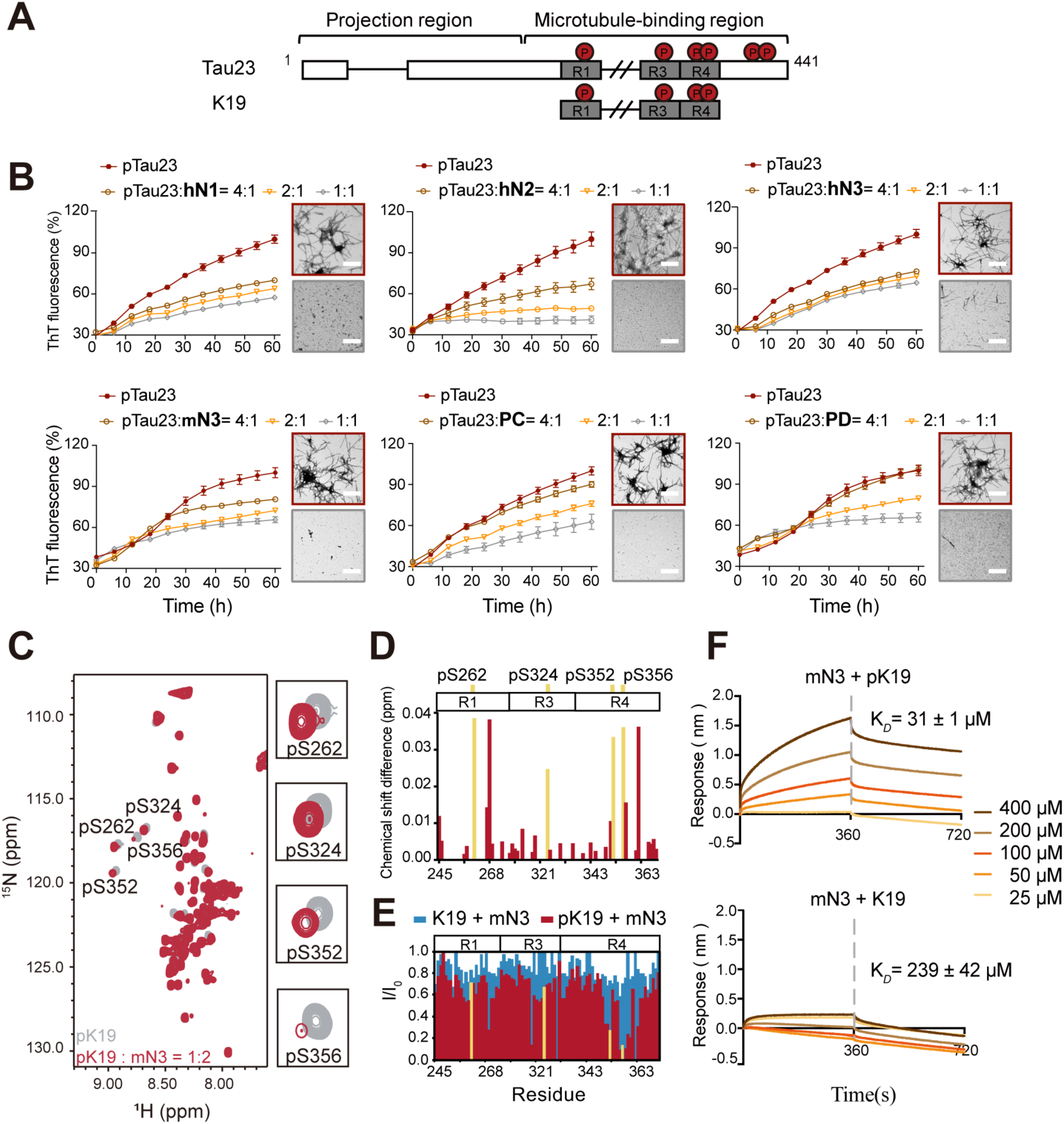
NMNATs inhibit pTau amyloid fibril formation and specifically bind to the phosphorylated sites of pTau. A Domain compositions of Tau23 and K19. The repeat regions are presented as gray boxes. Phosphorylation sites characterized in this work are marked. B Inhibition of NMNATs on the amyloid fibril formation of pTau23 (60 µM) measured by the ThT fluorescence kinetic assay. A gradient concentration of NMNATs was applied as indicated. The data shown correspond to mean ± s.d., with n = 5. C Overlay of 2D ^1^H-^15^N HSQC spectra of pK19 alone (100 µM, gray) and pK19 titrated by mN3 (200 µM, red). Signals of pSer residues are enlarged. D Residue-specific chemical shift changes of pK19 analyzed based on (c). The domain organization of pK19 is indicated and pSer residues are highlighted. E Residue-specific intensity changes of pK19 (red) and K19 signals (light blue) based on (c) and Supplementary Fig. 6a. F Binding affinity of pK19/K19 with mN3 measured by BLI. The association and dissociation profiles of pK19/ K19 to mN3 (20 µg ml^-1^) divided by a vertical dash line are shown. Concentrations of K19 proteins and dissociation constant (K_*D*_) are indicated.

To comprehensively investigate the activity of NMNAT on pTau aggregation, we next prepared different isoforms of NMNAT proteins from different organisms, including three isoforms of human NMNATs (hN1, hN2, and hN3), two isoforms of Drosophila NMNATs (PD and PC) and mouse NMNAT3 (mN3). We first confirmed that the NMNAT proteins contained normal enzyme activities **(Figure supplement 2)**. Then, we conducted the ThT fluorescence kinetic assay and transmission electron microscopy (TEM) to monitor their influences on the amyloid aggregation of pTau. The result showed that different NMNAT isoforms generally exhibited a potent chaperone activity against the amyloid aggregation of both pTau23 **(Figure 1B and figure supplement 3)** and pK19 **(Figure supplement 4)** in a dose-dependent manner. The results demonstrate that the chaperone activity of NMNAT against pTau aggregation is highly conserved in the NMNAT family across different organisms. Thus, NMNAT may serve as a molecular chaperone that is directly involved in the protection of pTau from pathological deposition.

### Mechanism of the interaction between mN3 and pTau

We next sought to investigate the structural basis of the interaction between NMNAT and pTau. Although hN2 represents the most biological relevant isoform to chaperone pathological pTau, purified hN2 is unstable and prone to aggregate *in vitro*, which hinders further structural characterization. Alternatively, we used mN3, which is of high thermostability and a highly conserved sequence to hN2 **(Figure supplement 5)**. We performed solution NMR spectroscopy and used mN3 to titrate ^15^N-labeled pK19. The 2D ^1^H-^15^N HSQC spectra showed a significant overall signal broadening of pK19, which indicates a strong interaction between mN3 and pK19 **(Figure 1C)**. In particular, the four phosphorylated Ser (pSer) residues showed large signal attenuations and chemical shift changes upon mN3 titration **(Figure 1D and 1E)**. Residues adjacent to pSer also exhibited prominent signal attenuations **(Figure 1E)**. Especially, repeat sequence R4 that contains two pSer residues showed the largest signal attenuations with I/I_0_ < 0.3 **(Figure 1E)**. In contrast, as we titrated non-phosphorylated K19 at the same ratio, only slight overall signal broadening was observed **(Figure 1E and figure supplement 6A)**. Moreover, pTau23, but not Tau23, showed significant chemical shift changes and intensity attenuations mainly on and around the pSer residues upon mN3 titration **(Figure supplement 6B)**. These results indicate that the pSer resides of pTau are the primary binding sites of mN3.

Further, to quantitatively measure the binding affinity between NMNAT and pTau, we conducted BioLayer Interferometry (BLI) analysis that is a label-free technology for measuring biomolecular interactions (Rich & Myszka, 2007). We immobilized mN3 on the biosensor tip and profiled the association and dissociation curves in the presence of either pTau or Tau **(Figure 1F and figure supplement 7)**. As we measured, the dissociation constant (*K*_*D*_) of the interaction between mN3 and pK19 is ∼ 31 µM, and that of mN3 and pTau23 is ∼ 9.9 µM **(Figure 1F)**. In contrast, the *K*_*D*_ of mN3 and K19 is ∼ 239 µM, and that of mN3 and Tau23 is ∼ 58.6 µM, which are about 10-fold weaker than that of the phosphorylated counterparts **(Figure 1F and figure supplement 7)**.

Taken together, these results indicate that MARK2-phosphorylation can significantly enhance the interaction between NMNAT and pTau through the specific interaction between NMNAT and the phosphorylated residues of pTau.

### mN3 utilizes its enzymatic substrate-binding site to bind pTau

To identify the binding site of mN3 for pTau, we firstly determined the atomic structure of mN3 at the resolution of 2.0 Å by X-ray crystallography **(Table supplement 1)**. mN3 formed a homo-dimer in the crystal packing with a buried surface area of 1,075.8 Å^2^ **(Figure supplement 8A)**, which is consistent with the molecular mass of a dimer (∼64 kDa) in solution measured by size exclusion chromatography and multi-angle laser light scattering (SEC-MALLS) **(Figure supplement 8B)**. The structure of mN3 protomer is similar to the known structure of hN3 with a r.m.s.d. value of 0.543 Å between Cα atoms **(Figure supplement 8C)**. The catalytic pocket that synthesizes NAD from NMN and ATP, is highly conserved in the NMNAT family **(Figure supplement 5 and 8C)**.

Next, we conducted a cross-linking mass spectrometry (CL-MS) with chemical cross-linker BS^3^ to covalently link paired lysine residues in spatial proximity (Cα-Cα distance < 24 Å) as pTau and mN3 interact, and then identified the cross-linked segments by mass spectrometry. The result showed 7 pairs of cross-linked segments between pK19 and mN3 with a confidence score of < 10^−7^ **(Table supplement 2)**. Lysine residues, K95, K139 and K206, that are involved in the cross-linking of mN3 with pK19, cluster around the entrance of the enzymatic pocket of mN3 (**Figure 2A**). The entrance of the enzymatic pocket feature a positively-charged patch mainly composed of residues K55, K56, R205 and K206 for the NMN and ATP binding **(Figure 2B and figure supplement 8C)**. This result implies that mN3 may utilize the same positively charged binding pocket for both pTau and the enzymatic substrates.

**Figure 2.**
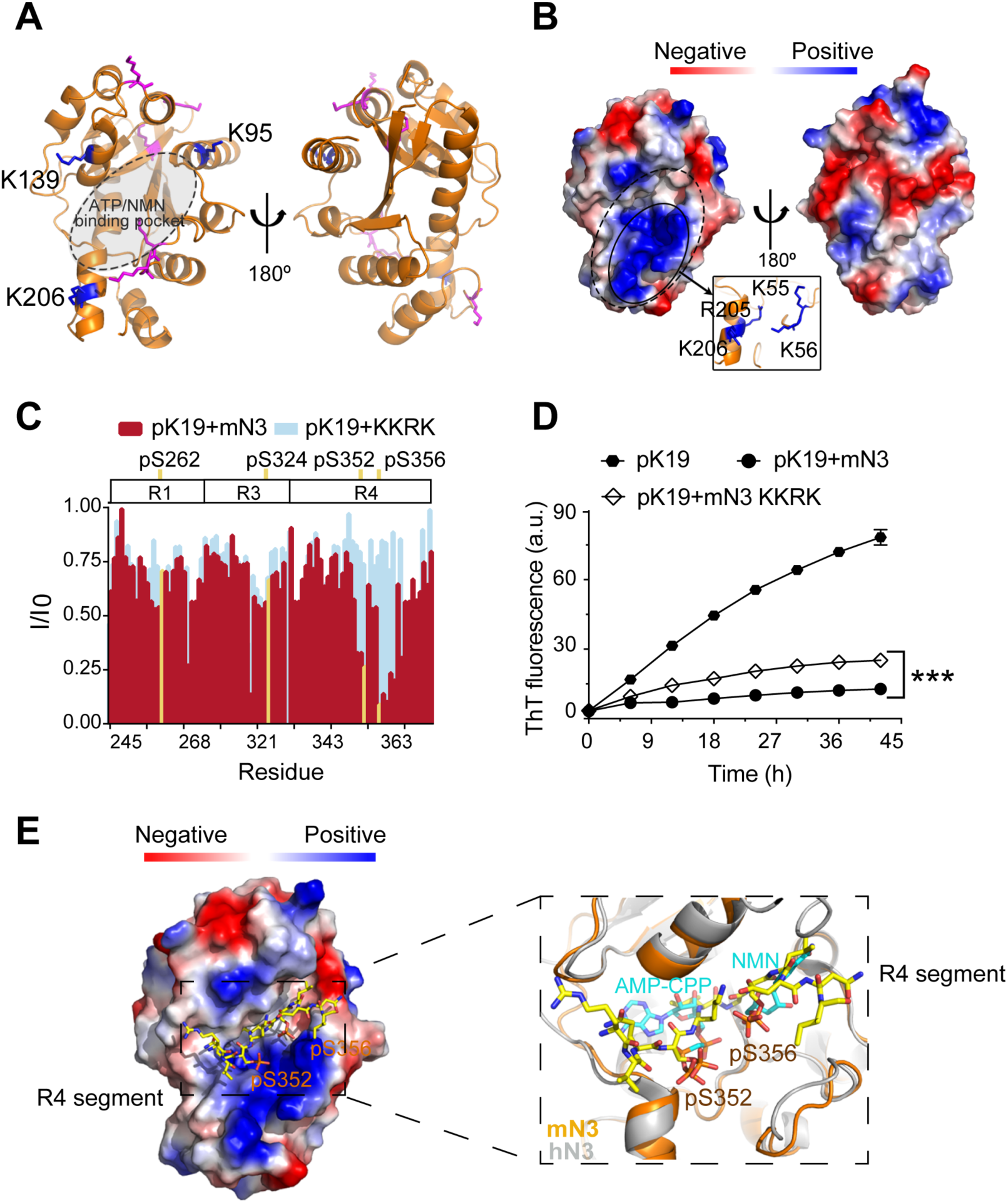
Structural characterization of pTau-binding site on NMNAT. A The structure of mN3 is shown in cartoon. Lysine resides that cross-linked with pK19 are shown as sticks in blue. Other lysine residues of mN3 are shown in magenta. The ATP/NMN binding site is shaded in gray. B Electrostatic surface representation of mN3. The positively charged patch of mN3 is indicated with solid ellipsoidal line. The residues that compose the positively charged patch are shown as sticks in blue in the zoom-in view. The hydrophobic area surrounding the positively-charged patch is indicated with dashed ellipsoidal line. C Overlay of the residue-specific intensity changes of pK19 signals titrated by mN3 (red) and KKRK mutant (light blue), respectively. pSer residues are colored in yellow. The domain organization of K19 is indicated on top. D Influence of KKRK mutations on the inhibition of mN3 against pK19 amyloid aggregation measured by the ThT fluorescence assays The molar ratio of pK19 to mN3 is 1:0.2. Data correspond to mean ± s.d., with n = 3. Values are compared using Student’s *t*-test. *\*\*\*P < 0.001*. E Structural model of mN3 in complex with the phosphorylated R4 segment ^349^ RVQ(p)SKIG(p)SLDNI^360^. The electrostatic surface of mN3 structure is shown. The peptide is shown as sticks in yellow. A zoom-in view of the peptide binding site in (E) superimposed on the structure of hN3 in complex with AMP-CPP and NMN (PDB ID: 1NUS). AMP-CPP and NMN are shown as sticks in cyan.

To further validate the pTau binding site on mN3, we constructed a quadruple mutation of K55E, K56E, R205E and K206E (referred to as mutation KKRK) to disrupt the positively charged interface. Differential scanning fluorimetry (DSF) confirmed that the mutations did not impair the overall structural stability of mN3 **(Figure supplement 9)**. To test whether the mutations influence the interaction between mN3 and pTau, we titrated pK19 with the KKRK mutant. The HSQC spectrum showed that the KKRK mutations significantly diminished the affinity of mN3 to pSer residues **(Figure 2C, figure supplement 10)**. Especially, the region (residues 349-360) of R4, that contains two pSer residues, exhibited a dramatically weakened binding to the KKRK mutant **(Figure 2C, figure supplement 10)**.

Furthermore, the KKRK mutations significantly impaired the chaperone activity of mN3 against the amyloid aggregation of both pK19 (**Figure 2D**) and pTau23 **(Figure supplement 11A)**. Note that the disruption of the positively charged patch did not completely eliminate the chaperone activity of mN3, indicating that other interactions also contribute to the binding of NMNAT to pTau. Of note, there is a hydrophobic area on the periphery of the positive-charge patch **(Figure 2B)**, implying that hydrophobic interactions may also contribute to the chaperone activity of MNNAT to pTau.

Since the NMR results identified the segment ^349^RVQ(p)SKIG(p)SLDNI^360^ shared by both pK19 and pTau23 as the primary binding segment of mN3, we next sought to characterize the complex structure of mN3 and pTau. We calculated and built the complex structure by Rosetta modeling **(Figure 2E)**. The complex structural model showed that the phosphorylated segment is well accommodated in the ATP/NMN-binding pocket of mN3. The phosphate groups of pS352 and pS356 orientate toward the positively charged pocket of mN3 and position in the same sites as those of ATP and NMN **(Figure 2E)**. This structure model explains the observation that mutations of K55, K56 and R205 (as in the KKRK mutant), which are the major components of the positively charged pocket and essential for the binding of phosphate groups, specifically abolished the binding of mN3 to the phosphorylated segment ^349^ RVQ(p)SKIG(p)SLDNI^360^ (**Figure 2C**). In addition, the chemical shift perturbations of pS262 of R1 and pS324 of R3 were also abolished as titrated by the KKRK mutant of mN3 **(Figure 2C)**, which indicates that these pSer residues may bind to mN3 in a similar manner as those of R4.

Primary sequence alignment shows that the key positively charged residues identified for pTau binding are highly conserved in the family of NMNATs from different species **(Figure supplement 5)**, which suggests that different NMNAT proteins employ a common and conserved interface for pTau binding. Indeed, mutation of the conserved K57 and R274 residues in hN2 severely impaired its chaperone activity against pTau23 aggregation **(Figure supplement 11B)**.

Taken together, these results indicate that NMNAT adopts a conserved pocket to bind both the enzymatic substrates and pTau. Thus, the binding of NMNAT to pTau is similar to the specific binding of enzyme and substrate.

### Competition between pTau and ATP/NMN for NMNAT binding

We next sought to understand how NMNAT spatially organizes its dual functions with the same pocket in a single domain. We have shown that the KKRK mutations of mN3 diminished the chaperone activity of NMNAT since it disrupts the positively charged pocket for the binding of phosphate groups. Next, we tested the influence of the KKRK mutations on the enzymatic activity of mN3 on NAD synthesis. The result showed that the KKRK mutations also eliminated the enzymatic activity of mN3, which is conceivable due to the inefficient binding of the mutant to the phosphate groups of ATP and NMN **(Figure 3A)**. On the other hand, we mutated H22, a key catalytic residue for NAD synthesis (Saridakis et al., 2001) that positions deep at the bottom of the substrate-binding pocket **(Figure supplement 12A)**. The result showed that the H22A mutation resulted in elimination of the enzymatic activity **(Figure 3A)**, which agrees with the previous study on the enzyme activity of NMNAT (Zhai et al., 2006). However, we found that the H22A mutation showed no influence on the chaperone activity of mN3 in inhibiting the amyloid aggregation of pK19 **(Figure 3B)**.

**Figure 3.**
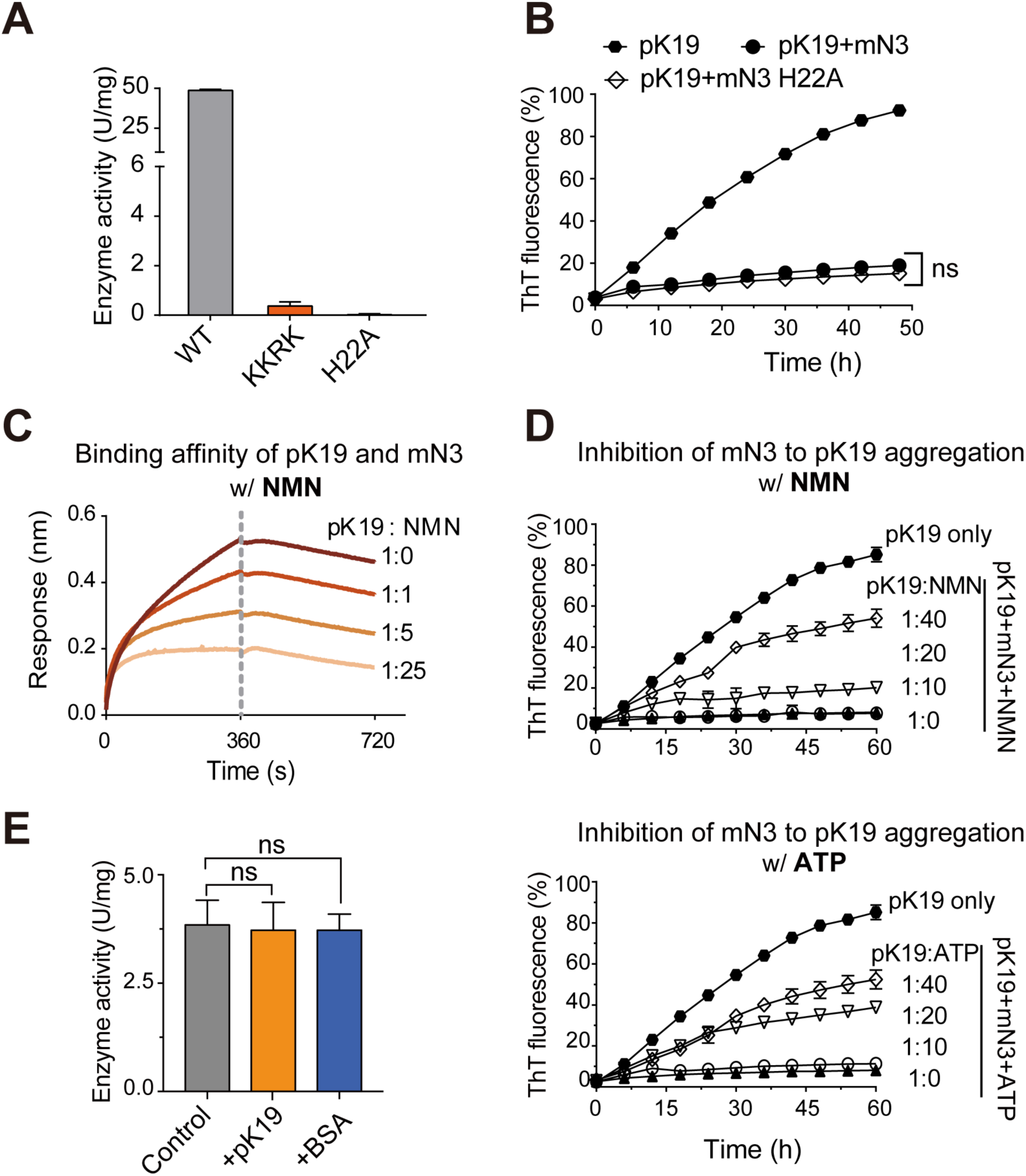
Competition of pK19 and NMN/ATP in the dual activities of NMNAT. A Enzyme activities of mN3 WT and mutants. The data shown correspond to mean ± s.d., with n = 3. B Influence of the H22A mutation on the inhibition of mN3 against the amyloid aggregation of pK19. The molar ratio of pK19 to mN3 is 1:0.2. The data shown correspond to mean ± s.d., with n = 3. Values are compared using Student’s *t*-test. “ns”, not significant C NMN weakens the binding of pK19 to mN3 in a dose-dependent manner measured by BLI analysis. The association and dissociation profiles are divided by a vertical dash line. The molar ratios of pK19/NMN are indicated. D The presence of NMN (top) or ATP (bottom) reduces the inhibitory effect of mN3 against pK19 amyloid aggregation in a dose-dependent manner. Molar ratios of pK19 to NMN/ATP are indicated. The data shown correspond to mean ± s.d., with n = 3. E The presence of pK19 (pK19: NMN/ATP = 10:1) shows no significant influence on the enzyme activity of mN3. The data shown correspond to mean ± s.d., with n = 3. Values are compared using Student’s *t*-test. “ns”, not significant

To examine the competition of the two activities, we used the BLI analysis and found that as the concentration of NMN increased, the binding of pK19 to mN3 was remarkably weakened **(Figure 3C)**. Consistently, the ThT assays showed that as the concentrations NMN or ATP decreased, the chaperone activity of mN3 on the amyloid aggregation of pK19 dramatically increased **(Figure 3D)**. In contrast, as we reversely added pK19 into the enzymatic reaction of NAD synthesis, no significant influence was observed **(Figure 3E)**.

Taken together, our data indicate that the enzymatic substrates (i.e. NMN and ATP) and the chaperone client pTau of mN3 share the same binding pocket with a partial overlap at the phosphate-binding site. While, ATP and NMN are superior to pTau on the mN3 binding.

### NMNAT protects pTau from aggregation and synaptopathy in *Drosophila*

To assess whether the specific binding of NMNAT to pTau characterized *in vitro* has functional relevance *in vivo*, we examined the protective capability of *Drosophila* isoform PD in a *Drosophila* tauopathy model. We overexpressed human wild type (Tau^WT^) or pathogenic Tau (Tau^R406W^) in the visual system of *Drosophila* using a photoreceptor-specific driver *GMR-GAL4* (Ali et al., 2012). The expression pattern can be easily visualized due to the highly organized parallel structure of the compound eye: the R1-R6 photoreceptors have their axons traverse the lamina cortex **(Figure 4A and 4B, magenta box)** and make synaptic contacts at the lamina neuropil **(Figure 4A and 4B, red box)**, while R7-R8 photoreceptors extend their axons beyond lamina and project to distinct layers in medullar neuropil **(Figure 4A and 4B, orange box)** (Sato, Suzuki, & Nakai, 2013).

**Figure 4.**
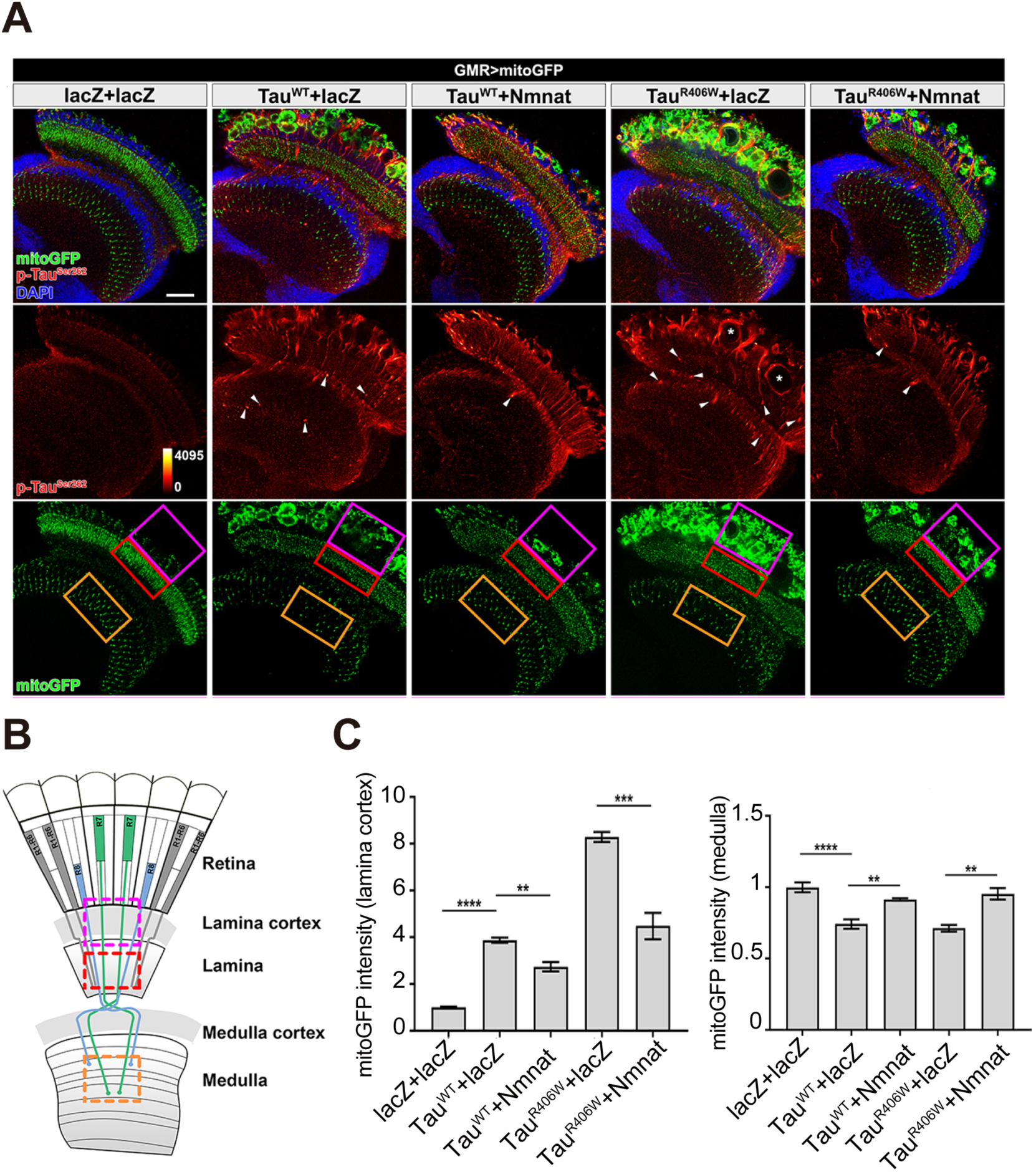
NMNAT (PD) suppresses pTau-induced mitochondrial clustering. A Adult female *Drosophila* (2 days after eclosion, DAE) brains expressing mitochondrial marker mitoGFP (Green) together with lacZ+lacZ, Tau^WT^+lacZ, Tau^WT^+NMNAT, Tau^R406W^+lacZ, or Tau^R406W^+NMNAT under photoreceptor specific driver *GMR-GAL4* were stained for pTau^Ser262^ (red spectrum) and DAPI (blue). White arrowheads show the aggregation of pTau. White asterisks show the holes formed in the lamina cortex layer, indicating retina degeneration. Magenta, red, and yellow boxes indicate lamina cortex, lamina, and medulla layers, respectively. Scale bar, 30 µm. B Diagram of the adult *Drosophila* visual system. Each ommatidia contains six outer photoreceptors (R1-R6) and two inner photoreceptors (R7 and R8). R1-R6 traverse the lamina cortex (magenta box) and project their axons into the lamina (red box), while the axons of R7 and R8 pass through the lamina and terminate in distinct synaptic layers in the medulla (orange box). C Quantification of mitoGFP intensity in the lamina cortex and medulla. Data are presented as mean ± s.d., with n =5. One-way ANOVA post hoc Tukey test; *\*\*P < 0.01, ***P < 0.001, ****P < 0.0001*.

As shown in **Figure 4A**, aggregation of pTau probed by antibody specifically recognizing pTau^Ser262^ was found in both Tau^WT^ and Tau^R406W^ fly brain, which was efficiently suppressed by PD overexpression **(Figure 4A and figure supplement 13)**. pTau aggregation impaired synaptic functions via (1) impairing mitochondrial dynamics and localization in neurons **(Figure 4C)**, (2) disrupting synaptic active zone integrity with misdistributed Bruchpilot (Brp) **(Figure supplement 14A and 14B)**, and (3) stimulating F-actin accumulation that restrict synaptic vesicle mobility and release **(Figure supplement 14C and 14D)**. These pTau pathology were remarkably enhanced in Tau^R406W^ fly. Importantly, overexpression of PD significantly suppressed pTau aggregation and protected against pTau-induced synaptic dysfunction mentioned above by restoring mitochondria and Brp localization at synaptic terminals, and alleviating pathological F-actin accumulation.

### mN3 mediates the recognition of Hsp90 to pTau to Hsp90

Previous studies showed that hN2 and Hsp90 co-precipitate with pTau in the brains of AD patients, and exhibit a synergistic effect in the attenuation of pTau pathology in cell models (Ali et al., 2012), Here, we used the SMPull (single-molecule pull-down) assay to identify the interplay between NMNAT, Hsp90 and pTau. SMPull is a powerful tool to quantitatively detect weak and transient interactions between protein complexes. As shown in **Figure 5A**, His_6_-tagged Hsp90 was coated on the slide, and the binding of pTau23 can be detected by the fluorescence from Alexa-647-labeled pTau23 using the total internal reflection fluorescence (TIRF) microscopy. The result showed that in the absence of mN3, binding of pTau23 to Hsp90 was only at the basal level that is similar to that of the blank slide (∼ 20 fluorescent spots per imaging area), which indicates that the interaction between them is very weak and transient. While, the addition of mN3 to the Hsp90/pTau23 system significantly increased the fluorescent spots in a dose-dependent manner (**Figure 5B, figure supplement 15A**). In contrast, the binding of non-phosphorylated Tau23 to Hsp90 is not affected by the addition of mN3 **(Figure supplement 15A and 15B)**. Thus, these results indicate that mN3 mediate the binding of pTau23 to Hsp90.

**Figure 5.**
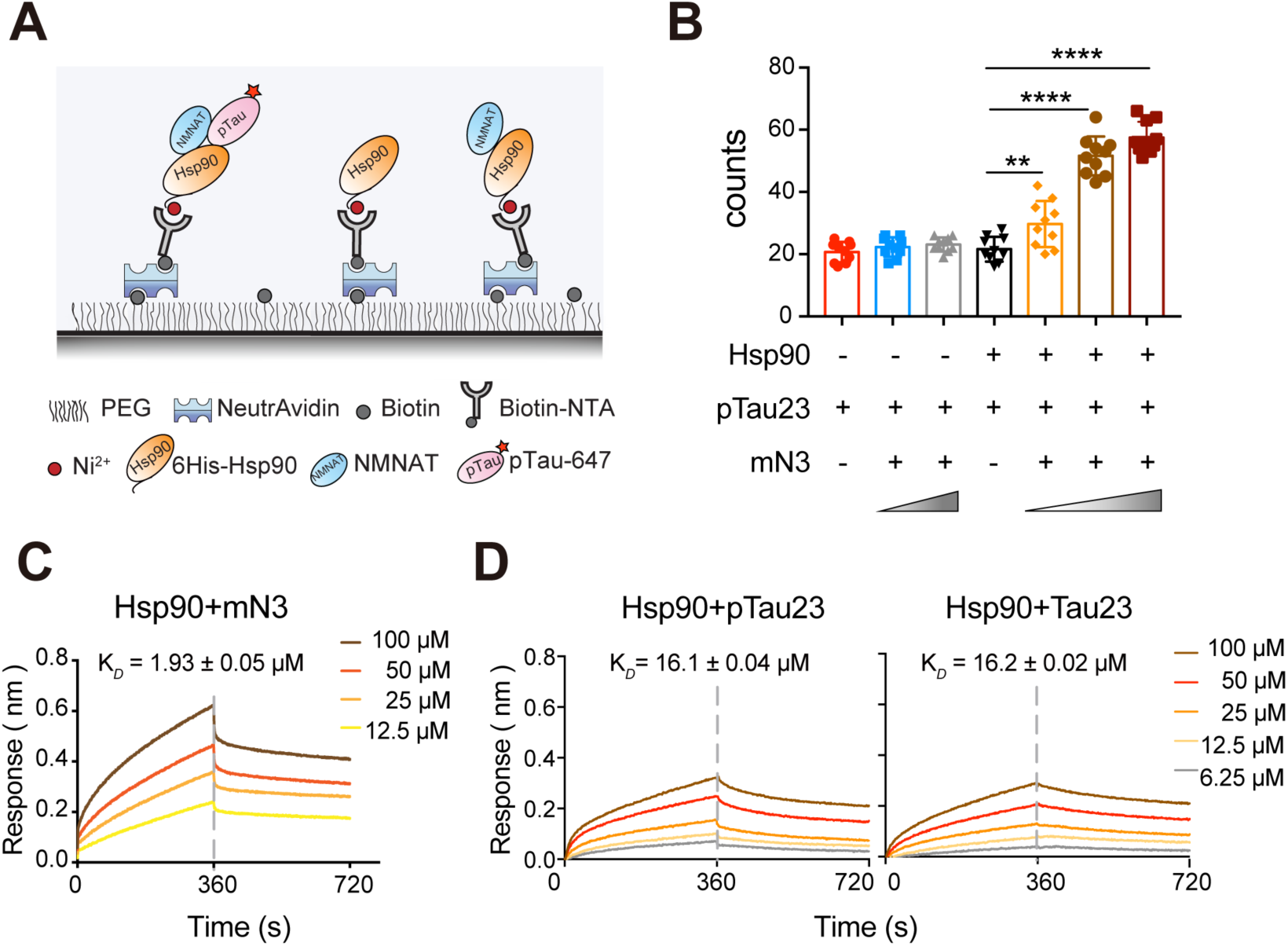
mN3 acts as a co-chaperone to assist Hsp90 in the recognition of pTau. A Schematic illustration of the SMPull assay by TIRF microscopy. His_6_-tagged Hsp90 was immobilized to the slide by chelating to Biotin-NTA-Ni. Single molecular interaction was monitored by the fluorescence from alexa-647 that was labeled on pTau23/Tau23. B The average number of fluorescent counts per imaging area detected by SMPull. TIRF images were recorded for the sample systems containing Hsp90, mN3 (4 nM) and pTau23 as indicated. The concentrations of mN3 from left to right are 0, 4, 20, 0, 0.8, 4, and 20 nM. Error bars denote standard deviations (s.d.) (n = 10). Values were compared using Student’s *t*-test. *\*\*, p < 0.01. ****, p < 0.0001*. C BLI measurements of mN3 binding to the SA sensor chip coated with biotinylated Hsp90 (20 µg ml^-1^). The mN3 concentrations are indicated. The K_*D*_ value of mN3 binding to Hsp90 is reported. D BLI measurements of the binding of pTau23 (left)/ Tau23 (right) to the SA sensor chip coated with biotinylated Hsp90 (20 µg ml^-1^). The Tau protein concentrations are indicated.

Furthermore, The BLI analysis showed that Hsp90 directly bound to mN3 with a K_*D*_ value of ∼ 1.93 µM (**Figure 5C**). While, Hsp90 was not able to differentiate pTau23 from Tau23 with the binding affinity of 16.1 µM to pTau23 and 16.2 µM to Tau23 **(Figure 5D)**. Taken together, our data indicate that NMNAT acts as a co-chaperone to assist Hsp90 in the recognition of of pTau.

## Discussion

### NMNAT represents a distinct class of molecular chaperones

NMNAT proteins have shown a robust neuroprotective activity in various models of neurodegenerative diseases correlated with the decrease of amyloid protein aggregation (Ali et al., 2013; Brazill, Li, Zhu, & Zhai, 2017; Conforti et al., 2014). In this work, we demonstrate that NMNAT functions as a molecular chaperone that selectively protect pTau from amyloid aggregation. Our work uncovers the structural basis of how NMNAT specifically binds pTau and how it manages its dual functions as both an enzyme and a chaperone. As illustrated in **Figure 6**, as NMNAT binds its enzymatic substrates, i.e. ATP and NMN, the substrates settle deep inside the pocket with defined interactions with NMNAT. As for the binding of pTau, the phosphorylated residues of pTau can specifically dock into the phosphate binding sites of NMNAT, which partially overlaps with the binding of ATP and NMN. Without phosphorylation, mN3 still show weak interactions with the KXGS motifs of Tau **(Figure 1E, figure supplement 6B)**, while MARK2-phosphorylation remarkably increases the binding affinity of mN3 to pTau by pSer specifically docking into the enzymatic pocket of mN3. Both KXGS motif recognition and phosphorylation might be required for the specific binding of mN3 to pTau.

**Figure 6.**
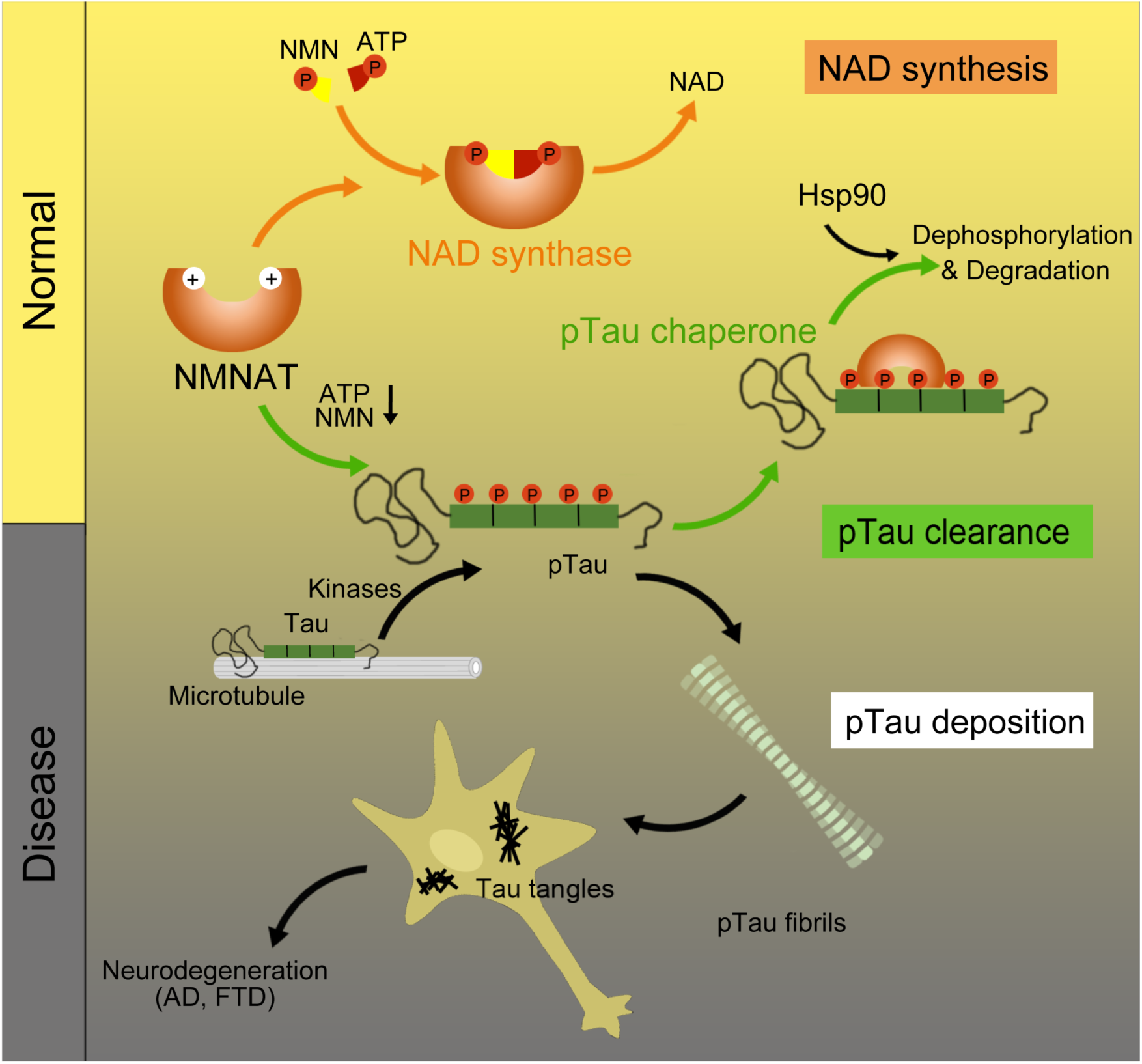
Schematics of NMNAT as a key node between pTau homeostasis and NAD metabolism. NMNAT functions as both an NAD synthase involved in NAD metabolism, and a molecular chaperone involved in the clearance of pathological pTau deposition. During aging as the level of ATP and NMN decrease, the chaperone function of NMNAT may show up to antagonize pTau aggregation.

Such specific binding features distinct NMNAT from other well-known chaperones. NMNAT is able to bind ATP as a substrate for its adenylyltransferase activity, while its chaperone activity is independent of ATP. NMNAT utilizes the same domain, even a shared pocket, for the binding of ATP and pTau, which explains the confusing previous observation that the neuroprotective effect of NMNAT is independent of ATP hydrolysis, yet requires the integrity of the ATP binding site(Ali et al., 2016). In addition, no large conformational change occurs as NMNAT binding to pTau.

In addition to inhibit pTau aggregation, NMNAT assists Hsp90 in the selection for pTau, implying that NMNAT serves as a co-chaperone of Hsp90 for pTau clearance. Previous studies identified CHIP protein as an important co-chaperone of Hsp90 for pTau removal (Dickey, Kamal, et al., 2007; Dickey, Patterson, et al., 2007). whereas the binding of CHIP to Tau is disrupted once Ser262/Ser356 of Tau is phosphorylated by MARK2 (Dickey, Kamal, et al., 2007). Therefore, Hsp90 may employ different cochaperones to handle different species of Tau and maintain Tau homeostasis.

### NMNAT links NAD and Tau metabolism

Our study demonstrates that the unique structural property of NMNAT gives rise to the binding of pTau with a high affinity comparable to that of an enzyme-substrate binding, but distinct from the weak interaction commonly found in chaperone-client binding (Koldewey, Horowitz, & Bardwell, 2017). The co-existence of the NAD synthase and chaperone activity sharing a common surface of NMNAT indicates an evolutionary connection between NAD metabolism and Tau proteostasis. During aging and AD pathogenesis, a concomitant decrease of ATP and NMN levels and pTau accumulation have been observed (Gomes et al., 2013; Stein & Imai, 2014; Wang & Mandelkow, 2015). Our data suggest that as ATP and NMN decrease, the shared enzymatic pocket may be more available for pTau binding. Therefore, the chaperone function of NMNAT may show up as ATP and NMN decrease during aging and neurodegeneration, implying a direct link between NAD metabolism and pTau proteostasis **(Figure 6)**. The cellular processes of NAD metabolism and Tau phosphorylation and proteostasis are complex with multiple nodes of regulation. NMNAT emerges as a critical regulator balancing the NAD-mediated active metabolic state and the amyloid-accumulating proteotoxic stress state. Such regulation would be particularly important for maintaining the structure and functional integrity of neurons.

## Acknowledgements

We thank Dr. Zhijun Liu, Dr. Songzi Jiang and other staff members of National Center for Protein science Shanghai for assistance in NMR data collection. This work was supported by the National Natural Science Foundation (NSF) of China (Grant No. 31470748), the Major State Basic Research Development Program (Grant No. 2016YFA0501902), the State High-Tech Development Plan (the “863 Program”) Award (Grant No. 2015AA020907), the “1000 Talents Plan” of China to C. Liu. The support also comes from Dr. John T. Macdonald Foundation (to C. Li.), the Lois Pope LIFE fellows Program (to. C. Li and Y. Zhu), NIH grant R56NS095893 (to R.G. Zhai), and Taishan Scholar Project (Shandong Province, People’s Republic of China) (to R.G.Z.).

## Materials and Methods

### Protein expression and purification

Genes encoding mN1, mN3, *Drosophila* PC, PD (gift from Dr. Yanshan Fang) and genes encoding hN1, hN3 (purchased from GENEWIZ, Inc. Suzhou, China) were amplified and inserted into pET-28a vector with an N-terminal His_6_-tag and a following thrombin cleavage site. Gene encoding hN2 (purchased from Genewiz, Inc.) was cloned into pET-32M-3C derived from pET-32a (Novagen). The resulting plasmid encodes a protein with an N-terminal MBP (maltose-binding protein) and a His_6_-tag followed by a HRV 3C protease recognition site. Mutations of mN3 including KK (K55EK56E), RK (R205EK206E), KKRK (K55EK56ER205EK206E) and H22A were constructed by site-directed mutagenesis using Q5^®^ Site-Directed Mutagenesis Kit (New England Biolabs). All the resulting constructs were verified by DNA sequencing (GENEWIZ, Inc. Suzhou, China).

NMNATs and variants were over-expressed in *E. coli* BL21 (DE3) cells. Cells were grown 2×YT medium at 37 °C to an OD_600_ of 0.8-1. Protein expression was induced by the addition of 0.2 mM isopropyl-β-d-1-thiogalactopyranoside (IPTG) and incubated at 16 °C for 15 hours. Cells were harvested by centrifugation at 4,000 *rpm* for 20 min and lysed in 50 ml lysis buffer (50 mM Tris-HCl, pH 8.0, 300 mM NaCl, and 2 mM phenylmethanesulfonyl fluoride (PMSF)) by a high-pressure homogenizer (800-1000 bar, 15 min). We next purified the over-expressed proteins by using HisTrap HP (5 ml) and HiLoad 16/600 Superdex 200 columns following the manufacturer’s instructions (GE Healthcare). The purified proteins were finally in a buffer of 50 mM Hepes-KOH, pH 8.0, 150 mM KCl, 10 mM MgCl_2,_ and 5% glycerol, concentrated, flash frozen in liquid nitrogen, and stored at −80 °C. The purity was assessed by SDS-PAGE. Protein concentration was determined by BCA assay (Thermo Fisher).

Human Tau23/K19 was expressed and purified on the basis of a previously described method (Barghorn, Biernat, & Mandelkow, 2005). Briefly, Tau23/K19 was purified by a HighTrap HP SP (5 ml) column (GE Healthcare), followed by a Superdex 75 gel filtration column (GE Healthcare). For ^15^N-or ^15^N/^13^C-labeled proteins, protein expression was the same as that for unlabeled proteins except that the cells were grown in M9 minimal medium with ^15^NH_4_Cl (1 g l^-1^) in the absence or presence of ^13^C_6_-glucose (2 g l^-1^).

### *In vitro* tau phosphorylation

Phosphorylation of Tau23/K19 by MARK2 kinase was carried out following a method described previously (Schwalbe et al., 2013). Breifly, Tau23/K19 was incubated with cat MARK2-T208E (a hyperactive variant) (Timm et al., 2003b) at a molar ratio of 10:1 in a buffer of 50 mM Hepes, pH 8.0, 150 mM KCl, 10 mM MgCl_2_, 5 mM ethylene glycol tetraacetic acid (EGTA), 1 mM PMSF, 1 mM dithiothreitol (DTT), 2 mM ATP (Sigma), and protease inhibitor cocktail (Roche) at 30 °C overnight. Phosphorylated Tau23/K19 was further purified by HPLC (Agilent) to remove kinase, and lyophilized. The sites and degrees of phosphorylation were quantified using 2D ^1^H-^15^N HSQC spectra according to previously published procedures (Eliezer et al., 2005; Schwalbe et al., 2013; Timm et al., 2003b)

### Thioflavin T (ThT) fluorescence assay

Amyloid fibril formation of pK19 and pTau23 were monitored using an *in situ* ThT-binding assay. The ThT kinetics for amyloid fibrils were recorded using a Varioskan Flash Spectral Scanning Multimode Reader (Thermo Fisher Scientific) with sealed 384-microwell plates (Greiner Bio-One). Client proteins were mixed in the absence or presence of NMNATs and variants in indicated molar ratios in a buffer of 50 mM Tris-HCl, 50 mM KCl, 5% glycerol, 0.05% NaN_3_, pH 8.0, respectively. A final concentration of 50 µM ThT was added to each sample. To promote the formation of amyloid fibrils, 5% (v/v) of fibril seeds (the seeds were prepared by sonicating fibrils for 15 s) were added to pK19 and pTau23, respectively. ThT fluorescence was measured in triplicates with shaking at 600 *rpm* at 37 °C with excitation at 440 nm and emission at 485 nm.

### Transmission electron microscopy (TEM)

5 µl of samples were applied to fresh glow-discharged 300-mesh copper carbon grids and stained with 3% v/v uranyl acetate. Specimens were examined by using Tecnai G2 Spirit TEM operated at an accelerating voltage of 120 kV. Images were recorded using a 4K × 4K charge-coupled device camera (BM-Eagle, FEI Tecnai).

### Nuclear magnetic resonance (NMR) spectroscopy

All NMR samples were prepared in a NMR buffer of 25 mM HEPES, 40 mM KCl, 10 mM MgCl2, and 10% (v/v) D2O at pH 7.0. NMR experiments were collected at 298 K on Bruker Avance 600 and 900 MHz spectrometers. Both spectrometers are equipped with a cryogenic TXI probe. Backbone assignments of K19 and pK19 were accomplished according to the previously published assignments (Eliezer et al., 2005) and validated by the collected 3D HNCACB and CBCACONH spectra, respectively. These experiments were performed using a ∼ 1 mM 15N/13C labeled sample. For HSQC titration experiments, each sample (500 µl) was made of 0.1 mM 15N labeled protein (K19/pK19/Tau/pTau), in the absence or presence of mN3 at a molar ration of 1:2. All NMR spectra were processed using NMRPipe ^64^ and analyzed using Sparky (Lee, Tonelli, & Markley, 2015) and NMRView (Johnson, 2004).

### Biolayer interferometry (BLI) assay

The binding affinity between mN3 and client proteins was inspected by BLI experiments with ForteBio Octet RED96 (Pall ForteBio LLC)(Rich & Myszka, 2007). All data were collected at 25 °C with orbital shaking at 1,000 rpm in 96-well black flat bottom plates (Greiner Bio-One). A total volume of 200 µl was used for each sample and all reagents were diluted in a buffer of 50 mM HEPES, 150 mM KCl, 10 mM MgCl_2_ at pH 8.0. Biotinylated mN3 or HSP90 (20 µg ml-1) was loaded onto SA sensors (ForteBio) for 180 s, followed by a 60 s baseline, and then associated with different concentrations of client proteins for 360 s. The association step was followed by a 360 s dissociation step. A reference sensor without client proteins was used to account for nonspecific binding of analyte to the sensor. All data were processed by data analysis software 9.0 (ForteBio). The competition of NMN with pK19 was monitored with the addition of different concentrations of NMN in both association and dissociation solution.

### mN3 crystallization, data collection and structure determination

Crystals of mN3 were obtained by the hanging drop vapor diffusion method at 18 °C. The condition of 0.04 M citric acid, 0.06 M Bis-Tris propane, pH 6.0-7.5, 20% PEG3350 yielded the diffraction quality crystals after 2 days. Before data collection, crystals were soaked in a cryoprotectant solution consisting of the reservoir solution and 10% (v/v) glycerol and then quickly frozen with liquid nitrogen.

Diffraction data of mN3 was collected at the wavelength of 0.9791 Å using an ADSC Quantum 315r detector at beamline BL17U of Shanghai Synchrotron Radiation Facility (SSRF). Diffraction data for the crystal was collected at 2.00 Å resolutions, as shown in Table 1. The intensity sets of the mN3 crystal was indexed, integrated and scaled with the HKL2000 package (Otwinowski & Minor, 1997).

The mN3 structure was solved by molecular replacement method using Phaser(McCoy et al., 2007) in the CCP4 crystallographic suite(Potterton et al., 2004) with the crystal structure of NMN/NaMN adenylyltransferase (1KQN) as template. Several cycles of refinement were carried out using Phenix and Coot(Emsley & Cowtan, 2004; Vagin et al., 2004) progress in the structural refinement was evaluated by the free R-factor.The mN3 structure belong to the P21 space group with cell dimensions a = 53.7 Å, b = 80.8 Å, c = 64.5Å.

### Size exclusion chromatography and multi-angle laser light scattering (SEC-MALLS)

The weight-average molecular weight (Mw) of mN3 was estimated by SEC-MALLS that consisted of a SEC column (KD–806 M, Shodex, Tokyo, Japan), a MALLS detector (DAWN HELEOS-II, = 658 nm, Wyatt Technologies, USA), and a RI detector (Optilab, = 658 nm, Wyatt Technologies, USA). 100ul mN3 (10 mg/ml) in a buffer of 50 mM Hepes-KOH, pH 8.0, 150 mM KCl, 10 mM MgCl_2,_ 5% glycerol and 0.05% NaN_3_ were loaded to the SEC column with a flow rate of 0.5ml/min at 25 °C. The resulting data was analyzed using ASTRA VI software software (Wyatt Technologies, USA).

### Cross-linking mass spectrometry analysis (XL-MS)

Cross-linking experiments were performed as described previously (Zhou et al., 2015). pK19 was incubated with mN3 at 6:1 molar ratio in a buffer containing 50 mM Hepes-KOH, 150 mM KCl at pH 8.0 for 20 min at 4 °C. Cross-linker BS^3^ (Thermo Fisher Scientific, 21585) was added at a 1:8 mass ratio and incubated at room temperature for 1 h. The reaction was quenched with 20 mM ammonium bicarbonate at room temperature for 20 min. Cross-linking products were analyzed by SDS-PAGE to assess the cross-linking efficiency. For MS analysis, proteins were precipitated with acetone; the pellet was resuspended in 8 M urea, 100 mM Tris (pH 8.5) and digested with trypsin at room temperature overnight. The resulting peptides were analyzed by online nanoflow liquid chromatography tandem mass spectrometry (LC-MS/MS). And the mass spectrometry data were analyzed by pLink(Yang et al., 2012).

### Differential scanning fluorimetry (DSF)

Thermal melting experiments were carried out using a QuantStudio™ 6 and 7 Flex Real-Time PCR Systems (Life) as described previously(Niesen, Berglund, & Vedadi, 2007). The buffer is 50mM HEPES, 150 mM KCl, 10 mM MgCl_2_, and 5% glycerol at pH 8.0. SYPRO Orange (Thermo Fisher) was added as a fluorescence probe at a dilution of 1:1000. 10 µL of protein mixed with SYPRO Orange (Thermo Fisher) solution (1:1000, 50mM HEPES, 150 mM KCl, 10 mM MgCl_2_, and 5% glycerol at pH 8.0) to a final concentration of 10 µM were assayed in 384-well plates (Life). Excitation and emission filters for SYPRO-Orange dye were 465 nm and 590 nm, respectively. The temperature was increased by 0.9 °C per minute from 25 °C to 96 °C. The inflection point of the transition curve (Tm) is calculated using protein thermal shift software v1.2 (Thermo Fisher).

### Modeling of the complex structure of peptide RVQ(p)SKIG(p)SLDNI and mN3

The 12-amino acid peptide ^349^RVQ(p)SKIG(p)SLDNI^360^ ((p)S: phosphoserine) was docked into mN3 following the rosetta flexile peptide docking (FlexPepDock) protocol (Raveh, London, Zimmerman, & Schueler-Furman, 2011) in Rosetta software package (Leaver-Fay et al., 2011). Firstly, the 12-mer peptide mimic RVQEKIGELDNI, in which the two phosphoserines were replaced by two glutamates, was docked to the crystal structure of mN3 (PDB: 5Z9R). We performed docking simulations with the restrains of two phosphate binding sites identified in the crystal structure of human cytosolic NMN/NaMN adenylyltransferase (PDB code: 1NUS) (X. Zhang et al., 2003). The extended 12-mer peptide mimic was initially placed near the putative phosphate binding site. 5,000 models were generated by using FlexPepDock protocol to simultaneously fold and dock the peptide over the receptor surface. In this fold-and-dock step, we imposed the distance restraints to confine the glutamate residues of peptide mimic within the phosphate binding sites identified from the crystal structure. The top models with favorable Rosetta energies and satisfied constrains were selected, and the phosphoserines were modeled back by replacing two phosphoserine mimic residues glutamates. The newly modeled structure was further refined by energy minimization to get rid of potential clash and maintain the identified phosphate binding site. After refinement, the top models ranked by Rosetta energies and constraints were selected for visually inspection.

### Enzyme activity assay

Enzyme activity of NMNAT was measured in a continuous spectrophotometric coupled assay by monitoring the increase in absorbance of NADH at 340 nm, The reaction process is as follows (Balducci et al., 1995):

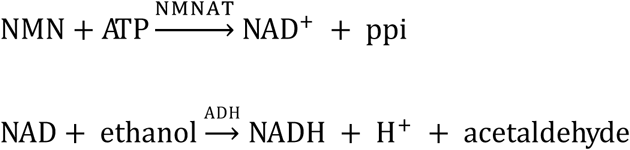

The reaction solution contains 28 mM HEPES buffer (pH 7.4), 11.2 mM MgCl2, 16 nM semicarbacide-HCl, 0.046 mM ethanol, 1.5 mM ATP, and 0.03 mg/ml yeast alcohol dehydrogenase (Sigma, A7011), and NMNAT or variants. The reaction was initiated by adding NMN to a final concentration of 0.625 mM. All measurements were performed at 37 °C. The activity was calculated using the equation below.

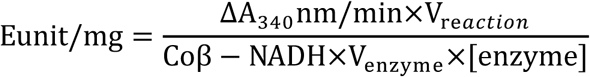

Where C_0_β-NADH, the extinction coefficient of β-NADH at 340 nm, is 6.22 (Zhai et al., 2006).

### *Drosophila* stocks and genetics

Flies were maintained on cornmeal-molasses-yeast medium at 25 °C, 65% humidity, 12 h light/dark cycle. The following strains were used in this study: *UAS-Tau*^*WT*^ and *UAS-Tau*^*R406W*^ obtained from Dr. Mel Feany(Ali et al., 2012), *UAS-mitoGFP* obtained from Dr. Hugo J. Bellen (Duncan Neurological Research Institute, Baylor College of Medicine), *GMR-GAL4* and *UAS-LifeAct-GFP* obtained from Bloomington Stock Center.

### Immunohistochemical staining of fly brains

Fly brains with attached lamina were dissected and stained as previous described (Li et al., 2017). Briefly, flies were dissected in cold PBS (pH 7.4) and fixed in freshly made 4% formaldehyde for 15 min. Brains were washed in PBS containing 0.4% (v/v) Triton X-100 (PBTX) and incubated with primary antibodies at 4 °C overnight. Brains were then washed with PBTX and incubated with secondary antibodies at room temperature for 2 h. After that, brains were stained with 4’,6-diamidino-2-phenylindole (DAPI; Invitrogen, Carlsbad, CA, USA) for 10 min and mounted with VECTASHIELD Antifade Mounting Medium (Vector Laboratories Inc., Burlingame, CA, USA). Samples were kept at 4 °C until imaging. The following antibodies were used in this study: anti-Brp (1:250, Developmental Studies Hybridoma Bank, East Iowa City, IA, USA) and anti-pTau^Ser262^ (1:250, Santa Cruz Biotechnology, CA, USA).

### Confocal imaging and processing

Fly brains were imaged using an Olympus IX81 confocal microscope coupled with a 60× oil immersion objective. Images were processed using FluoView 10-ASW software (Olympus) and analyzed using Fiji/Image J (version 1.52). The intensity data were plotted as mean ± SD and statistical analyses were performed using One-way ANOVA post hoc Tukey test by Graphpad Prism (version 7.04).

### Single-molecule imaging and spot counting

Purified pTau23/Tau23 proteins was mixed with a 3-fold Alexa Fluor™ 647 (Thermo Fisher, A32757) in a reaction buffer (50 mM NaH2PO4/Na2HPO4 at pH 7.4, 150 mM KCl, 0.5 mM TCEP) at 37 °C for 1 h. The labeled proteins were further purified using the Superdex 200 columns (GE Healthcare, USA) in a buffer containing 50mM NaH2PO4/Na2HPO4 at pH 7.4, 150 mM KCl, 0.5 mM TCEP.

All single-molecule assays were performed in the working buffer including 50 mM NaCl, 50 mM Tris, pH 8.0 and 0.1 mM TCEP at room temperature. Single-molecule imaging was conducted in the working buffer containing an oxygen scavenging system consisting of 0.8 mg/ml glucose oxidase, 0.625% glucose, 3 mM Trolox and 0.03 mg/ml catalase to minimize photobleaching. Slides were firstly coated with a mixture of 97% mPEG and 3% biotin-PEG, flow chambers were assembled using strips of double-sided tape and epoxy. Neutravidin and 20 nM biotin-NTA (Biotium) charged with NiCl2 were sequentially flowed into the flow chamber and each was incubated for 5 min in the working buffer. The immobilization of HSP90 (5 nM) was mediated by surfaced-bound Ni2+. Next, 4 nM Tau23/pTau23 and various concentrations of mN3 were added and incubated with the immobilized HSP90 for 10 min before data acquisition. An objective type total internal reflection fluorescence (TIRF) microscopy was used to acquire single-molecule data. Alexa647 labeled Tau23 or pTau23 was excited at 647 nm with a narrow band-pass filter (ET680/40 from Chroma Technology). Single-molecule analysis was performed using software smCamera. Mean spot per image (imaging area 2500 µm^2^) and standard deviation were calculated from 10 different regions.

---

**Supplementary Figure 1.**
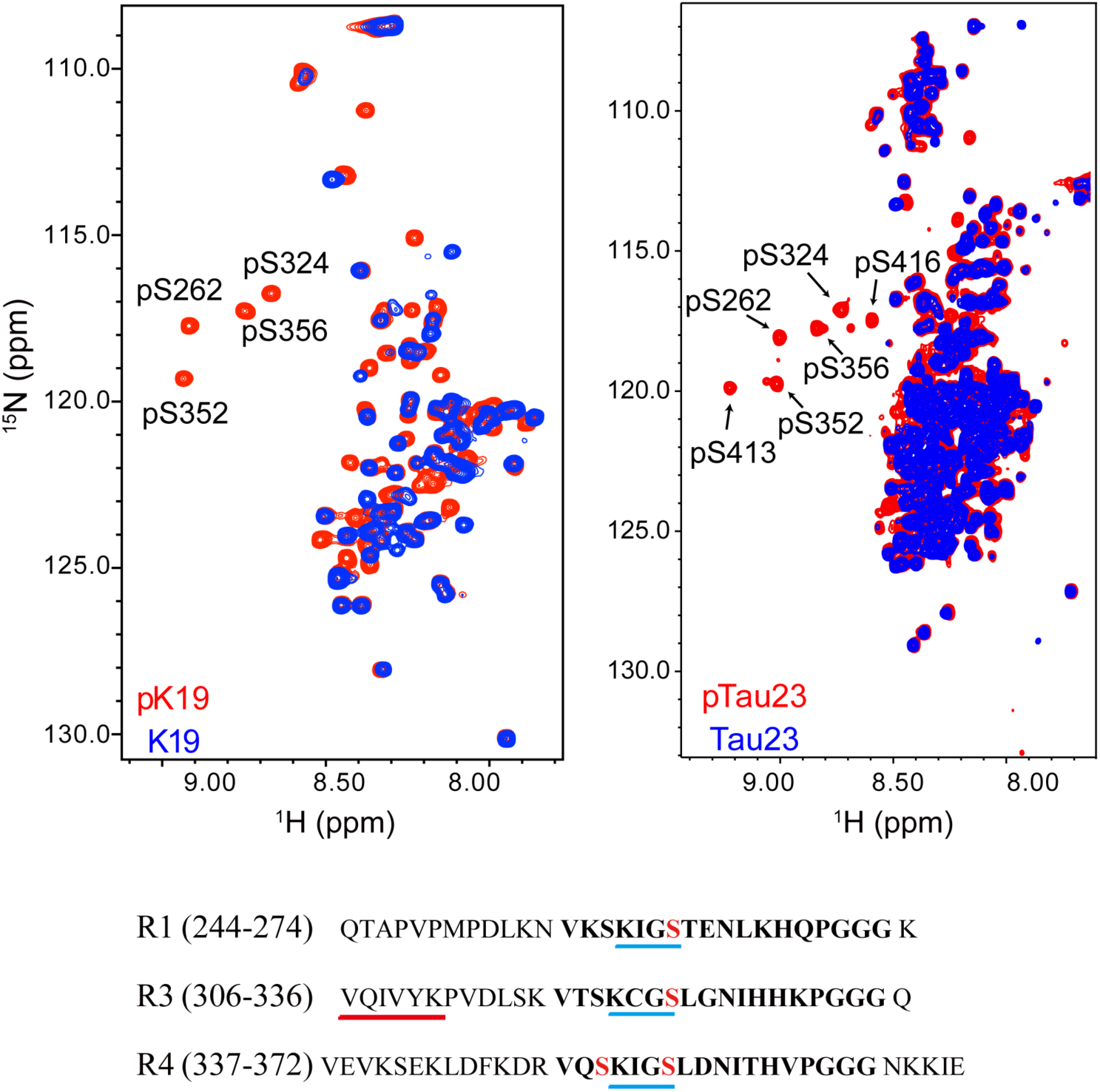
Characterization of MARK2 phosphorylation sites on pK19 (left) and pTau23 (right) by NMR. Overlay of the 2D ^1^H-^15^N HSQC spectra of pK19 (red) and K19 (blue), and pTau23 (red) and Tau23 (blue). The phosphorylated serine residues are labeled. The primary sequences of R1, R3 and R4 are shown. The 18-residue repeats are in bold. The KXGS motifs are underscored with blue lines. The four serine residues that are phosphorylated by MARK2 are highlighted in red. The fibril-forming core sequence is underscored with a red line.

**Supplementary Figure 2.**
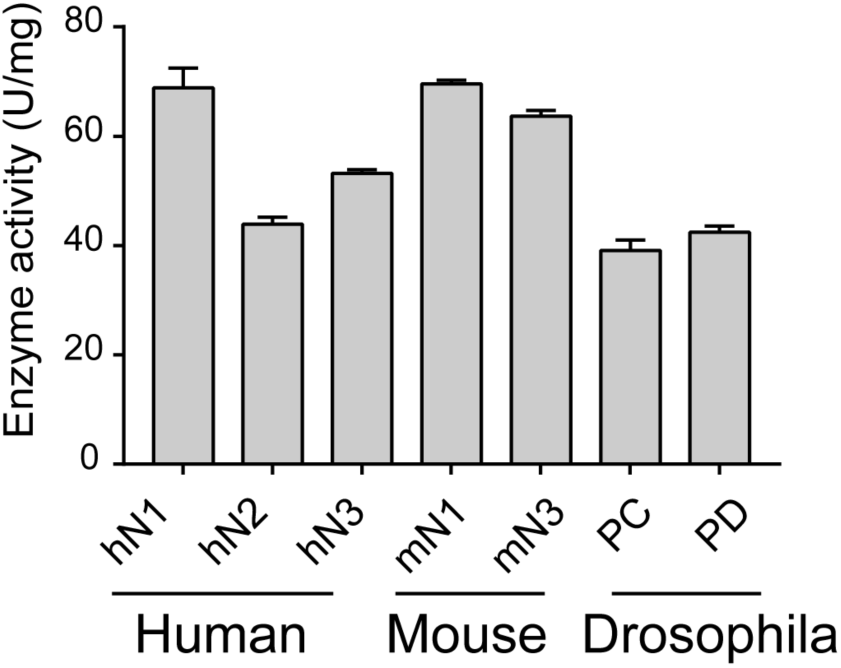
Enzyme activities of different isoforms of NMNAT from different organisms. The data shown correspond to mean ± s.d., with n = 3.

**Supplementary Figure 3.**
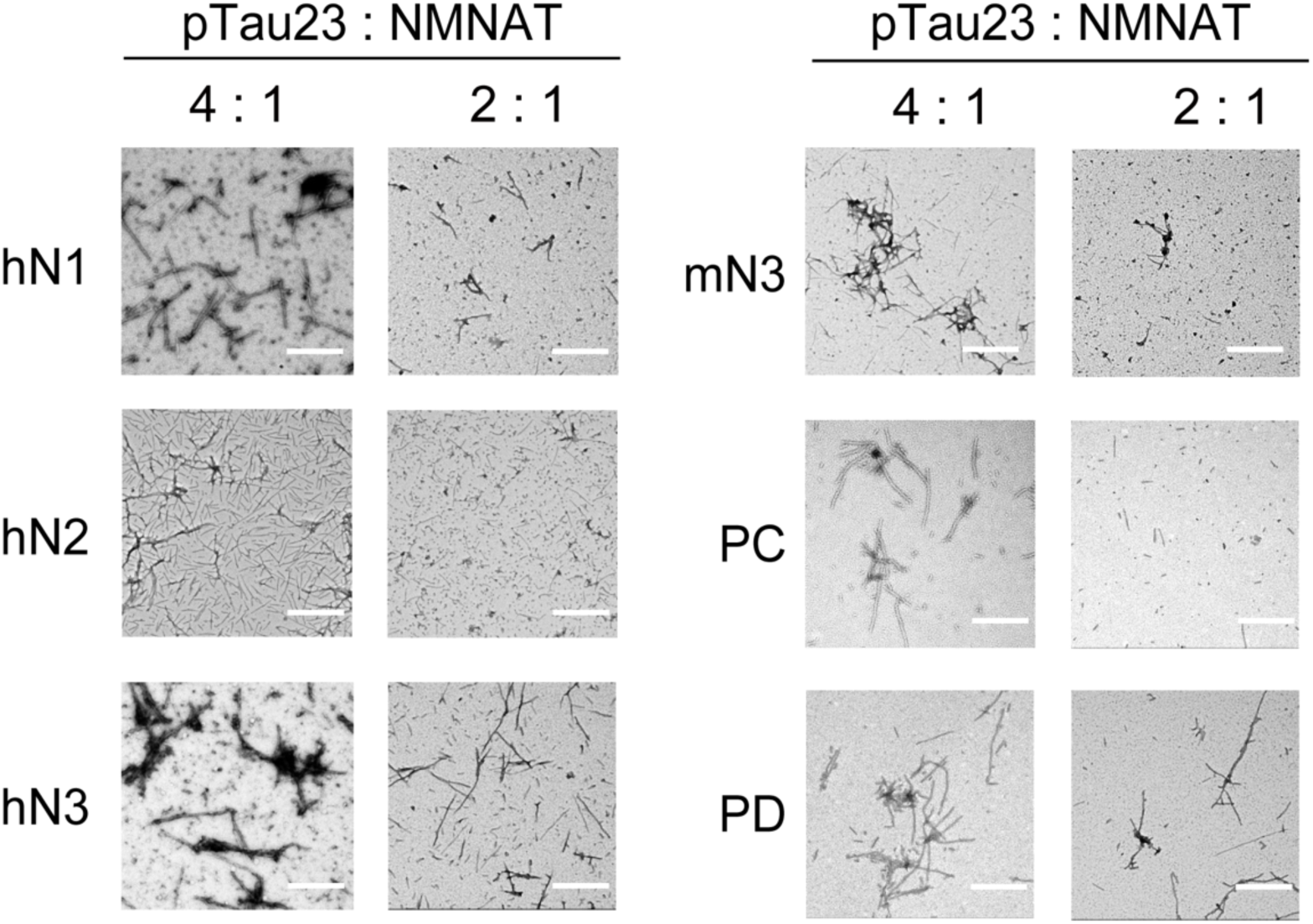
Inhibition of NMNATs on the fibril formation of pTau23 imaged by negative-staining TEM. Tau fibrils were imaged after pTau23 incubation with NMNATs at 37 °C for 60 hours. The molar ratios of pTau23 to NMNATs are indicated. Scale bars: 500 nm.

**Supplementary Figure 4.**
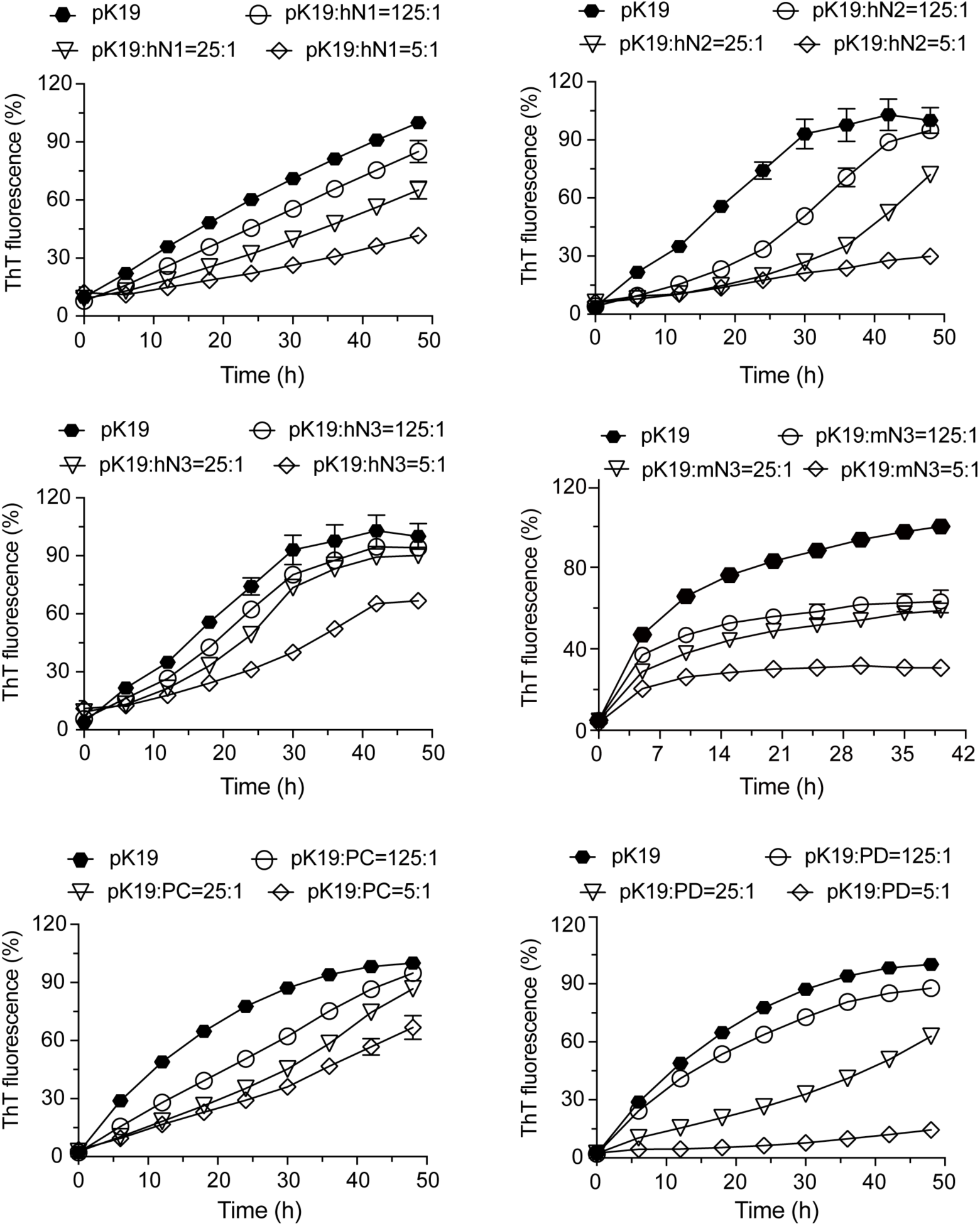
Inhibition of NMNATs on the fibril formation of pK19 measured by ThT kinetic assay. Inhibition of NMNATs on the amyloid fibril formation of pK19 (100 µM) measured by the ThT fluorescence kinetic assay. A gradient concentration of NMNATs was applied as indicated. The data shown correspond to mean ± s.d., with n = 3.

**Supplementary Figure 5.**
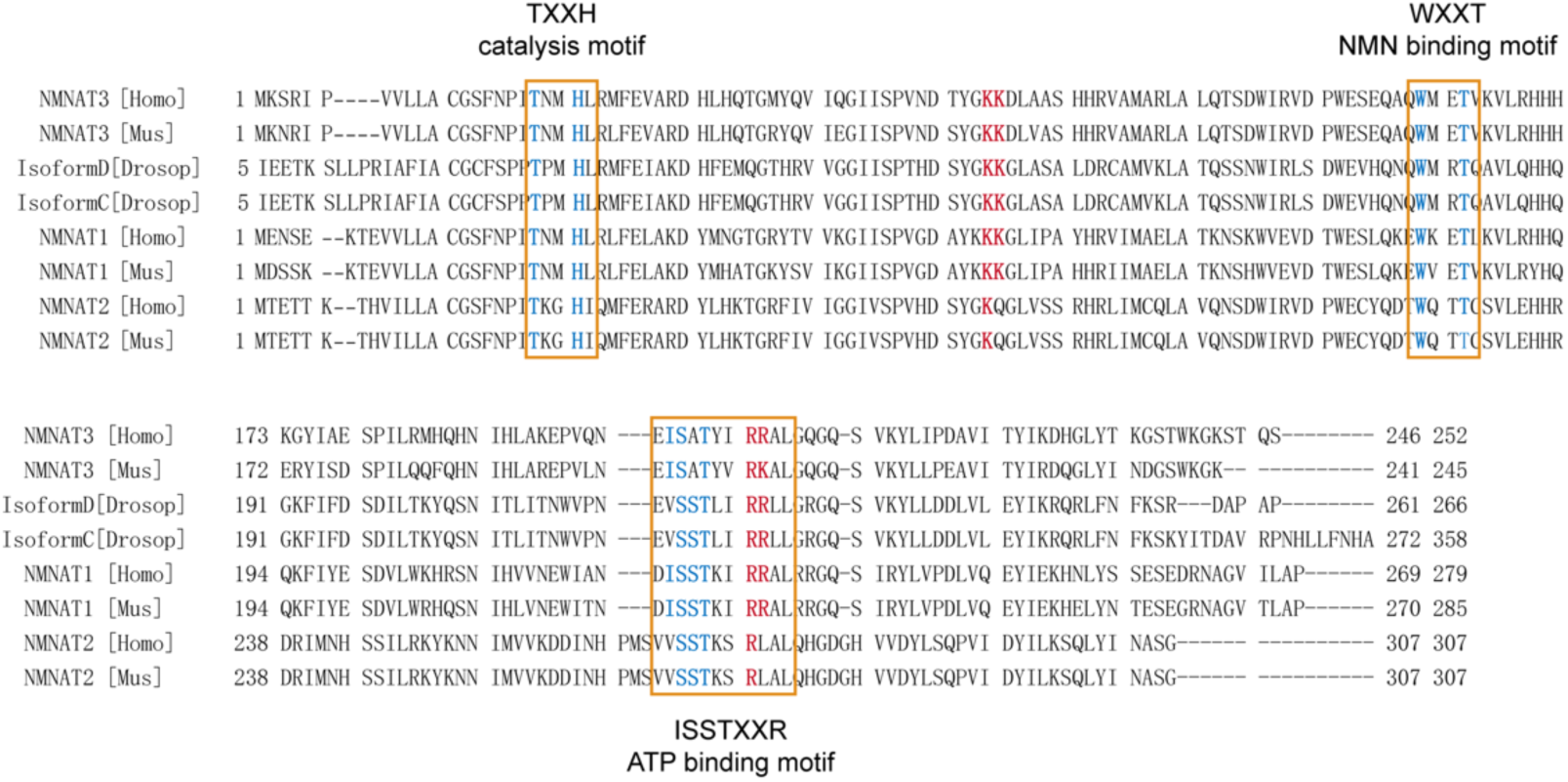
Sequence alignment of NMNATs from different species. The catalytic and substrate-binding motifs (highlighted with orange frames) are highly conserved and the conserved residues are in blue. The positively charged residues that are essential for the chaperone activity are colored in red.

**Supplementary Figure 6.**
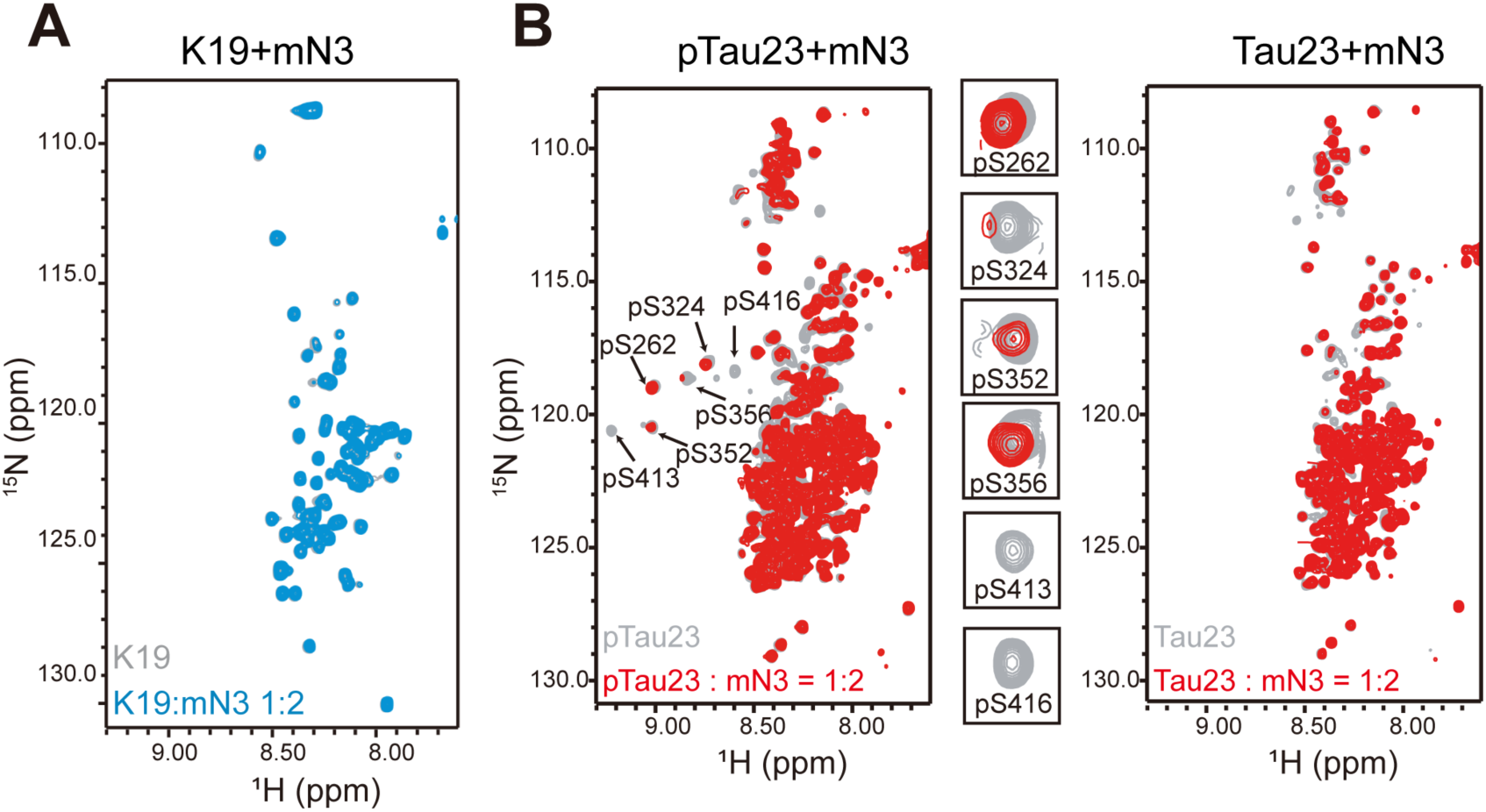
NMR titration of NMNAT to Tau. A Overlay of 2D ^1^H-^15^N HSQC spectra of K19 alone (100 µM, gray) and K19 titrated by mN3 (blue; molar ratio 1:2). B Overlay of 2D ^1^H-^15^N HSQC spectra of pTau23 (left)/Tau23(right) alone (100 µM, gray) and that titrated by mN3 (red; molar ratio 1:2). pSer residues are enlarged.

**Supplementary Figure 7.**
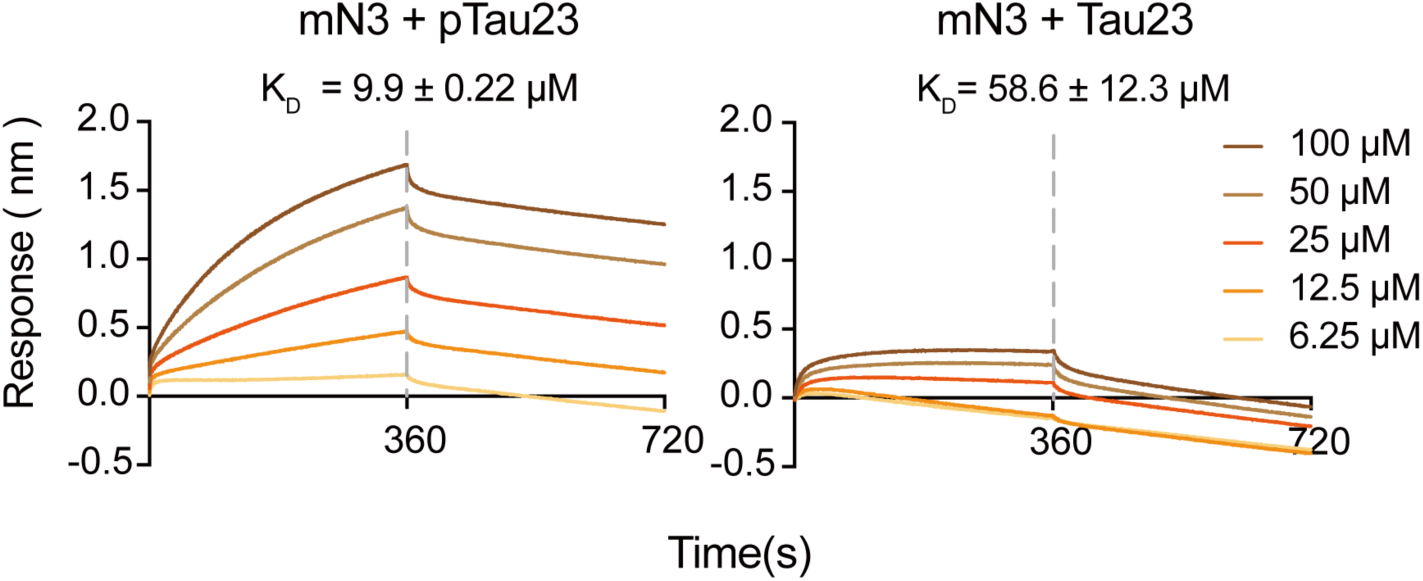
Binding affinity of pTau23 (left) and Tau23 (right) with mN3 measured by BLI. The SA sensor chip was coated with biotinylated mN3 (20 µg ml^-1^). The association and dissociation profiles divided by a vertical dash line are shown. The K_*D*_ values are reported.

**Supplementary Figure 8.**
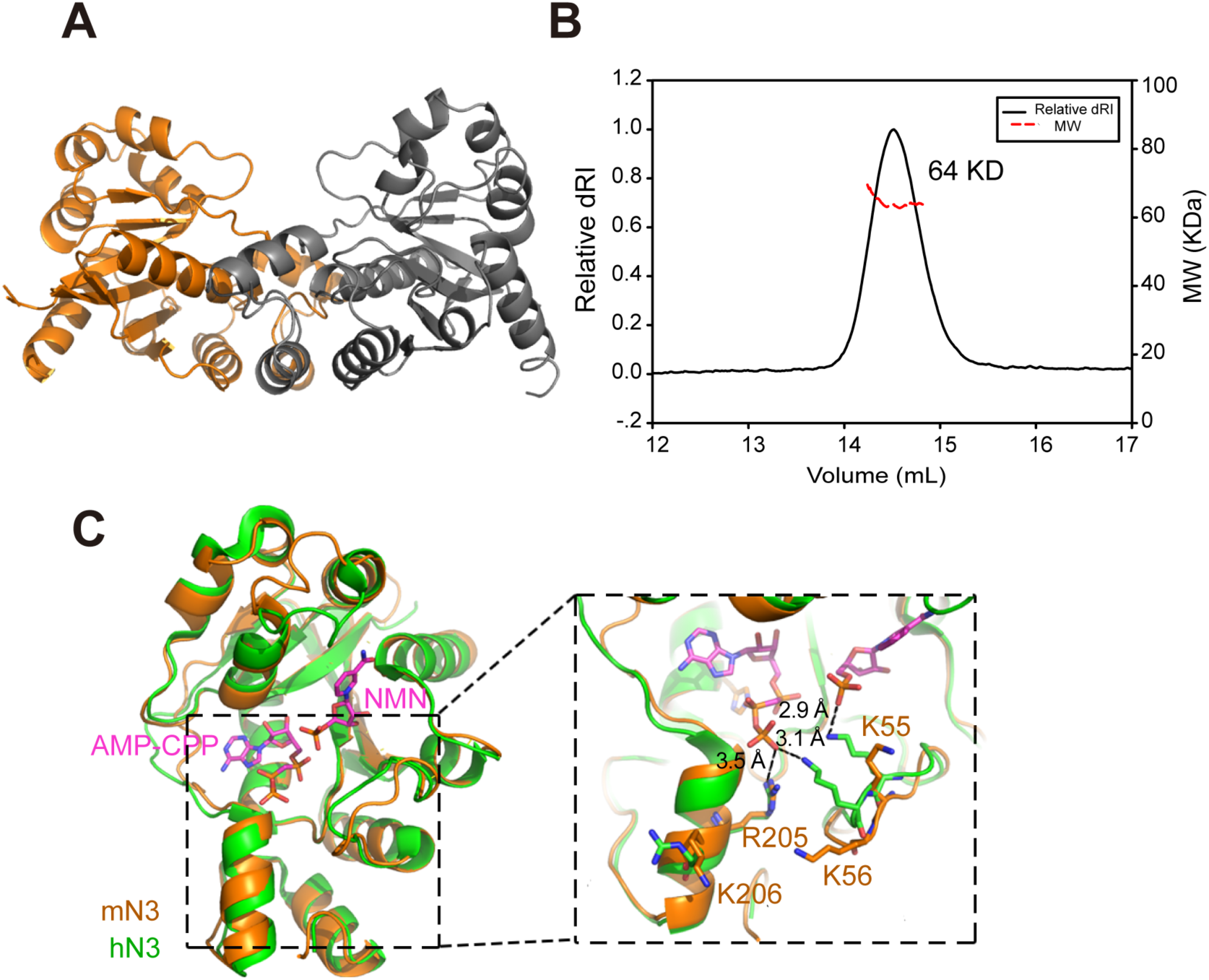
Structural characterization of mN3. A Crystal structure of mN3. Two protomers are colored in orange and dark gray, respectively. B The variations of differential refractive index (dRI, left y-axis) and molecular mass (right y-axis) are shown. The smooth trace is the elution profile of mN3 from size exclusion chromatograph. The red dotted line shows the resulting molecular mass (∼64 kDa), indicating a dimer of mN3 in solution. C Structure superimposition of mN3 (orange) and hN3 (green, PDB ID: 1NUS). Substrates AMP-CPP (non-hydrolysable ATP analogue) and NMN from the structure of hN3 are shown as sticks in magenta. Positively charged residues that compose the positively-charged patch of mN3 are shown in sticks. Their interactions with the phosphate groups of substrates are labeled in the zoom-in view.

**Supplementary Figure 9.**
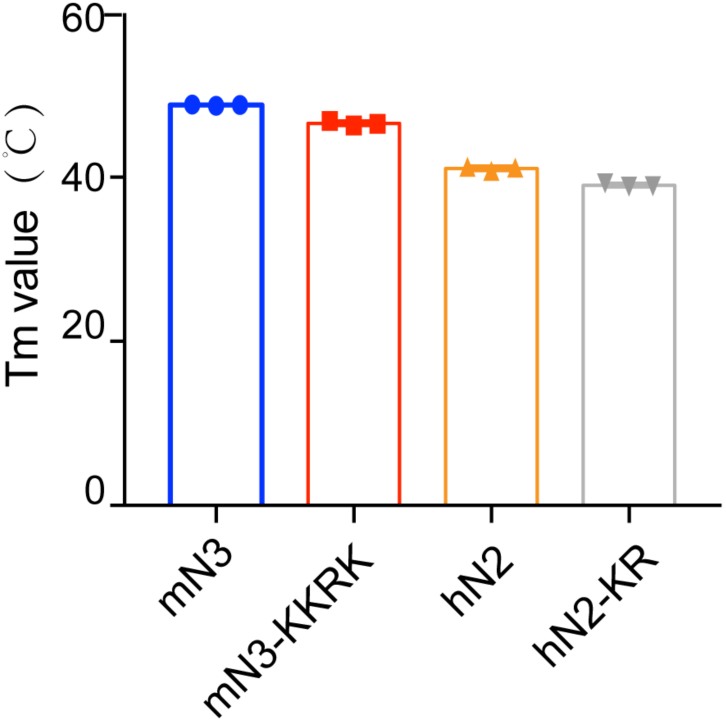
Melting temperature of mN3 and hN2 variants measured by the DSF assay. The data shown correspond to mean ± s.d., with n = 3.

**Supplementary Figure 10.**
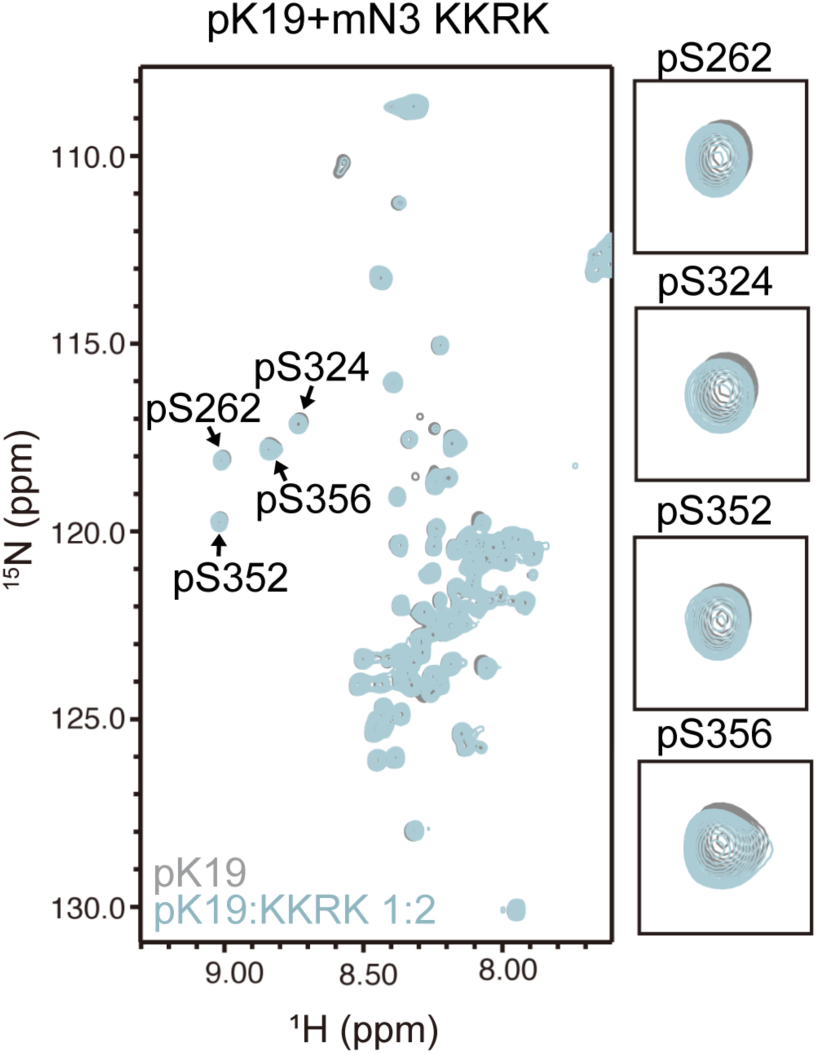
2D ^1^H-^15^N HSQC spectra of pK19 titrated by mN3-KKRK. pK19: 100 µM, gray; pK19 titrated by mN3-KKRK: molar ratio of 1:2, light blue. Signals of pSer residues are enlarged.

**Supplementary Figure 11.**
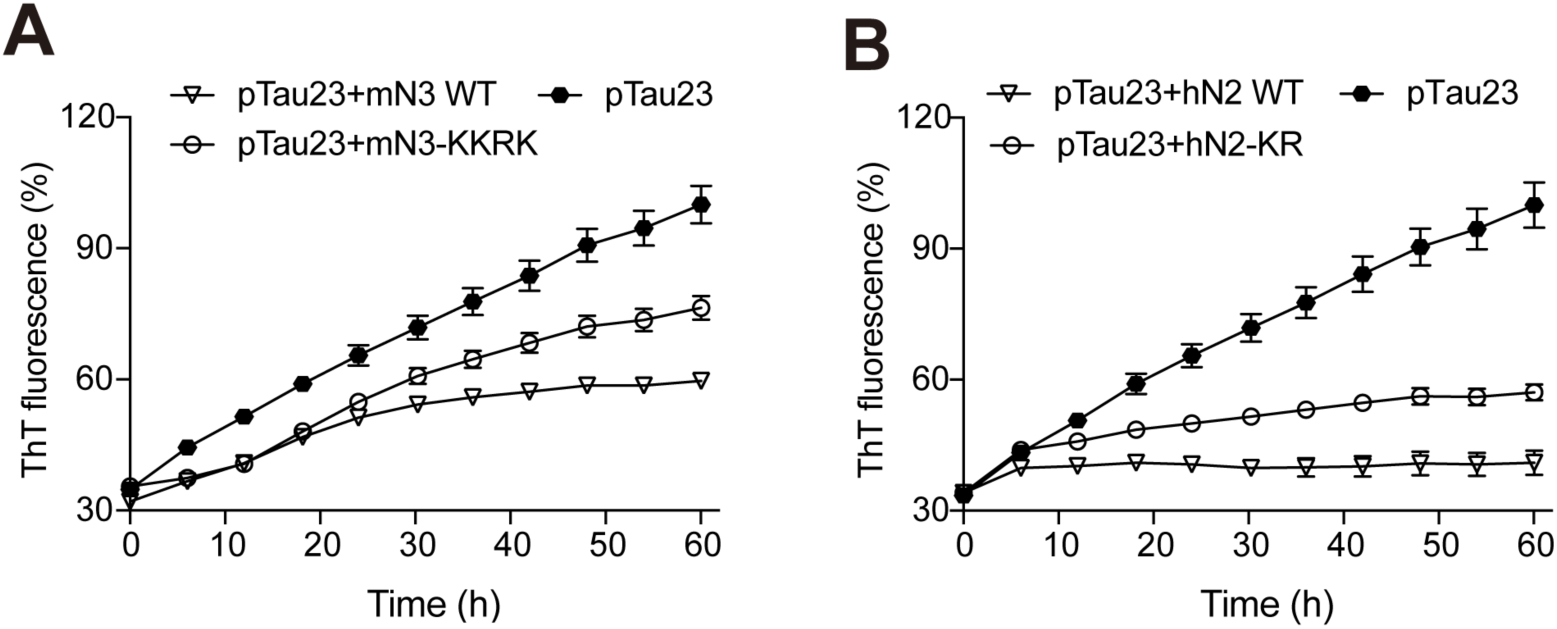
Influences of mN3 (A) and hN2 (B) mutations on pTau23 aggregation measured by ThT fluorescence assays. The molar ratios of mN3 and hN2 mutants to mN3 are 1:1, respectively. The data shown correspond to mean ± s.d., with n = 5.

**Supplementary Figure 12.**
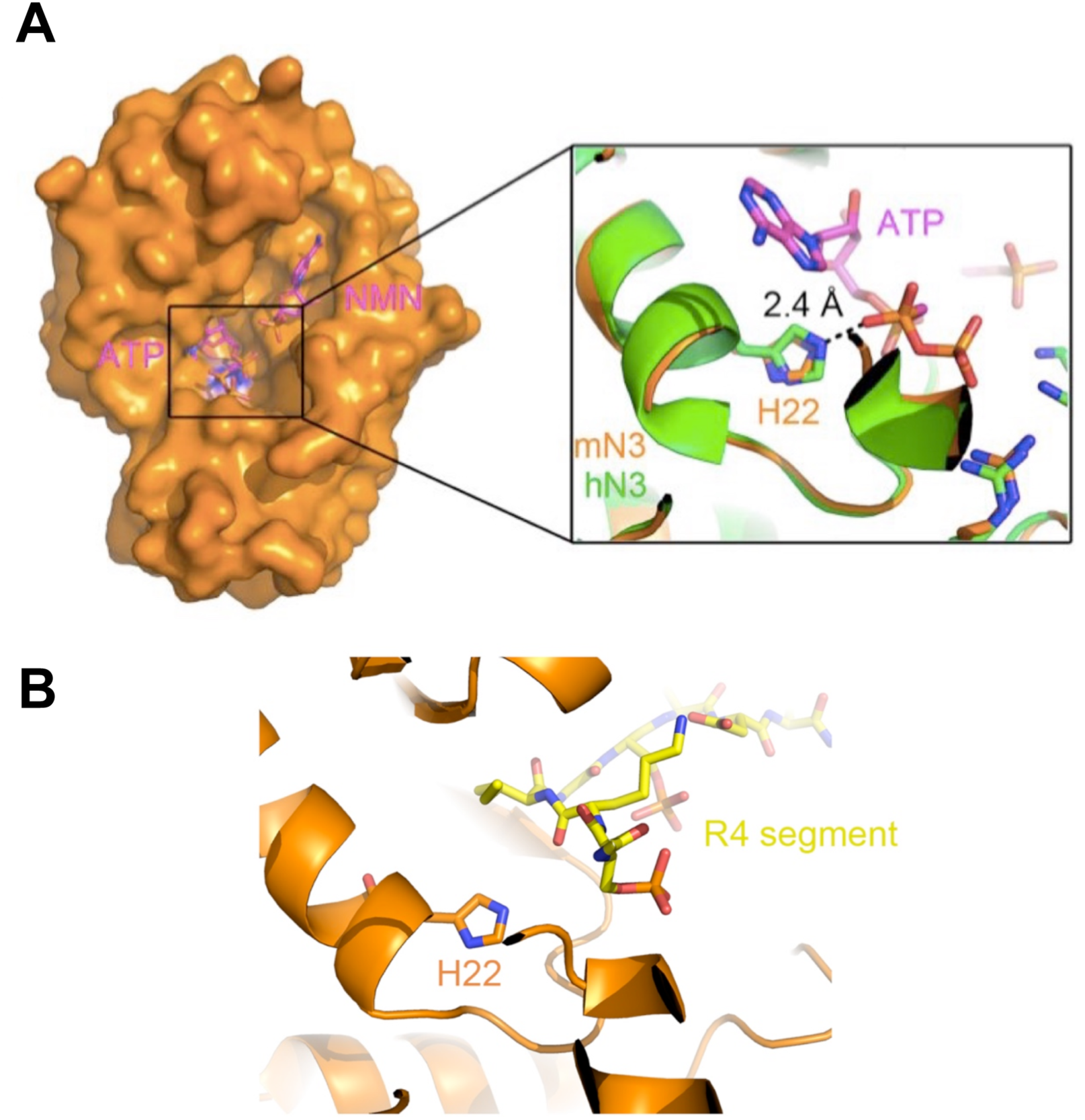
Position of H22 in the over-all mN3 structure. A The structure surface of mN3 is shown in orange. The apo structure of mN3 is superimposed with the structure of hN3 in complex with AMP-CPP and NMN (PDB ID: 1NUS). AMP-CPP and NMN are shown in magenta as sticks. The interaction of H22 and AMP-CPP is shown in a zoom-in view. B R4 segment resides at the entrance of the enzymatic pocket of mN3, and thus the mutation of of H22, which is deep at the bottom of the pocket, may not significantly influence the binding of R4 to mN3. R4 segment is in yellow sticks. mN3 is in orange.

**Supplementary Figure 13.**
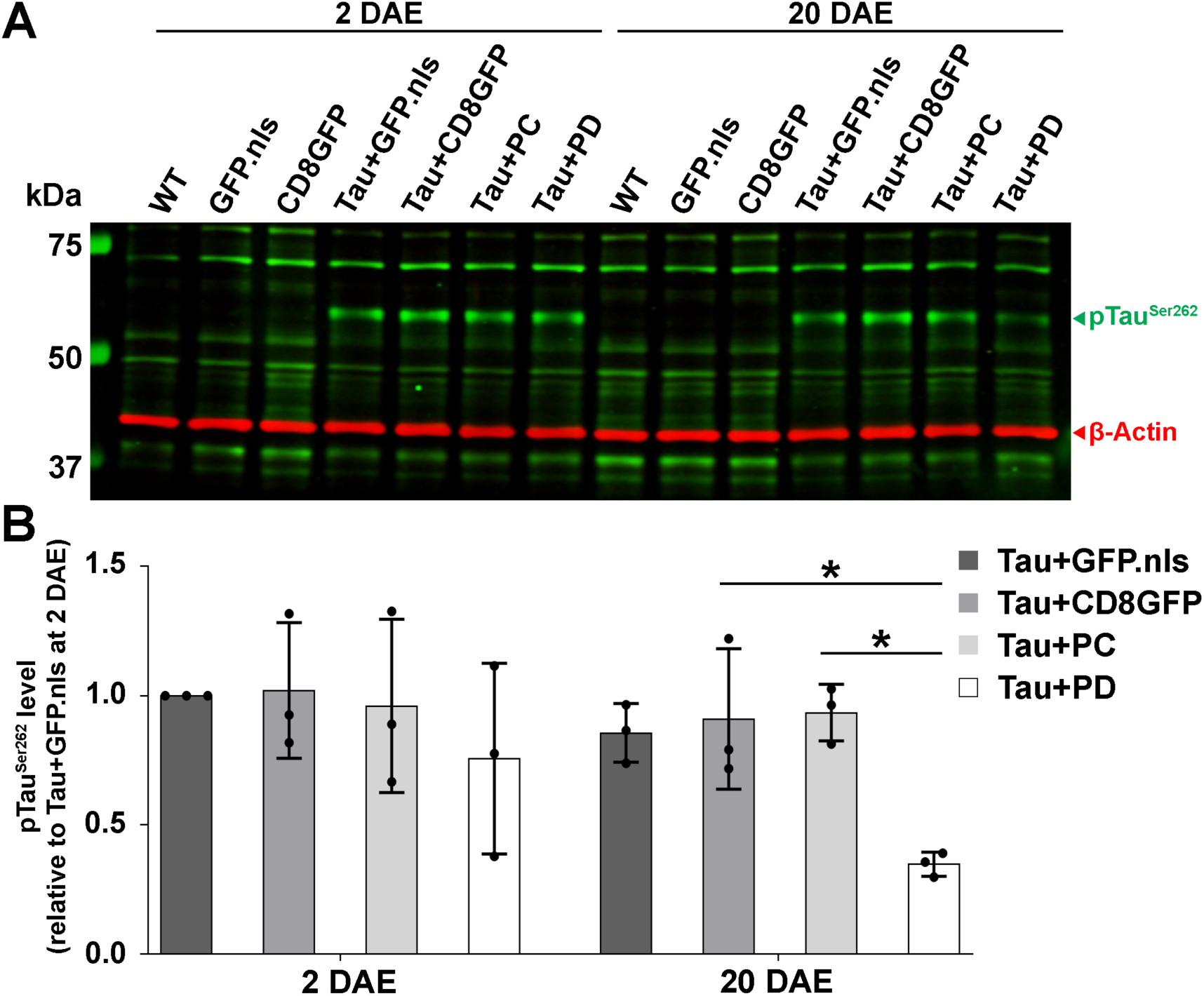
NMNAT reduces the levels of Ser262 hyper-phosphorylated Tau. D Brain lysates of 2 or 20 DAE wild-type flies or flies overexpressing GFP, hTau with GFP or NMNAT (PC PD) were probed with antibodies specific for disease-associated phospho-Tau epitopes Tau Ser262 and actin for loading control. E Quantification of pTau species normalized to actin and Tau+GFP from samples in (A) in 2 DAE and 20 DAE flies. *, P <0.05.

**Supplementary Figure 14.**
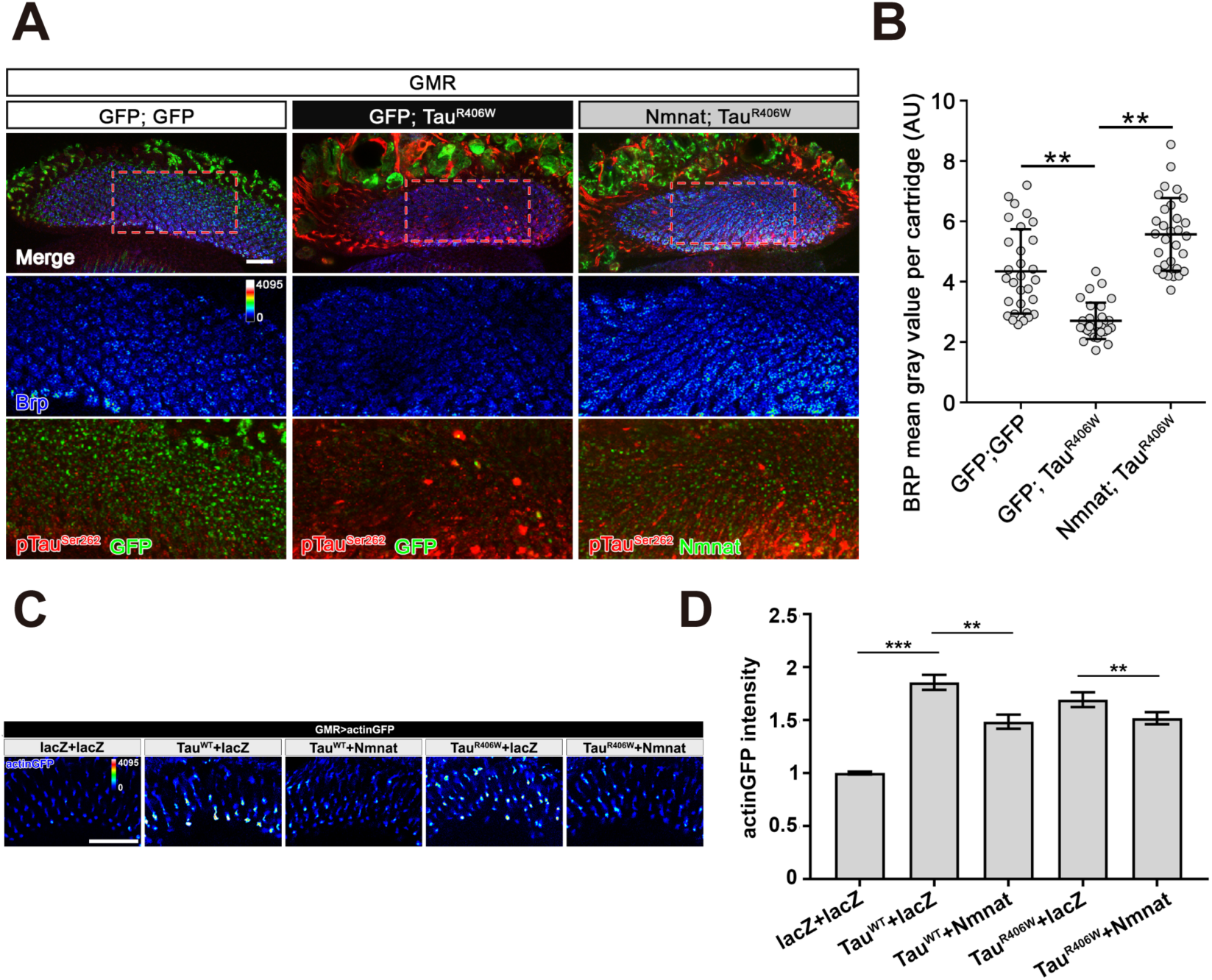
NMNAT (PD) suppresses pTau-induced Brp loss and F-actin accumulation at synaptic terminals. A Confocal micrographs show the Brp staining in lamina synapses. Brp labels active zone structure. pTau^Ser262^ labels hyperphosphorylated Tau. Arrowheads mark pTau aggregates. Scale bar, 20 µm. B Quantification (mean ± s.d.; n = 30 cartridges from three animals) of the Brp level in the lamina. One-way ANOVA post hoc Tukey test; \*\**P < 0.01*. C Medulla area of adult female *Drosophila* (2 DAE) expressing actin-GFP (blue spectrum) together with lacZ+lacZ, Tau^WT^+lacZ, Tau^WT^+NMNAT, Tau^R406W^+lacZ, or Tau^R406W^+NMNAT under photoreceptor specific driver *GMR-GAL4*. Scale bar, 30 µm. D Quantification (mean ± s.d.; n = 5) of the actin-GFP level in the medulla. One-way ANOVA post hoc Tukey test; *\*\*P < 0.01, ***P < 0.001*.

**Supplementary Figure 15.**
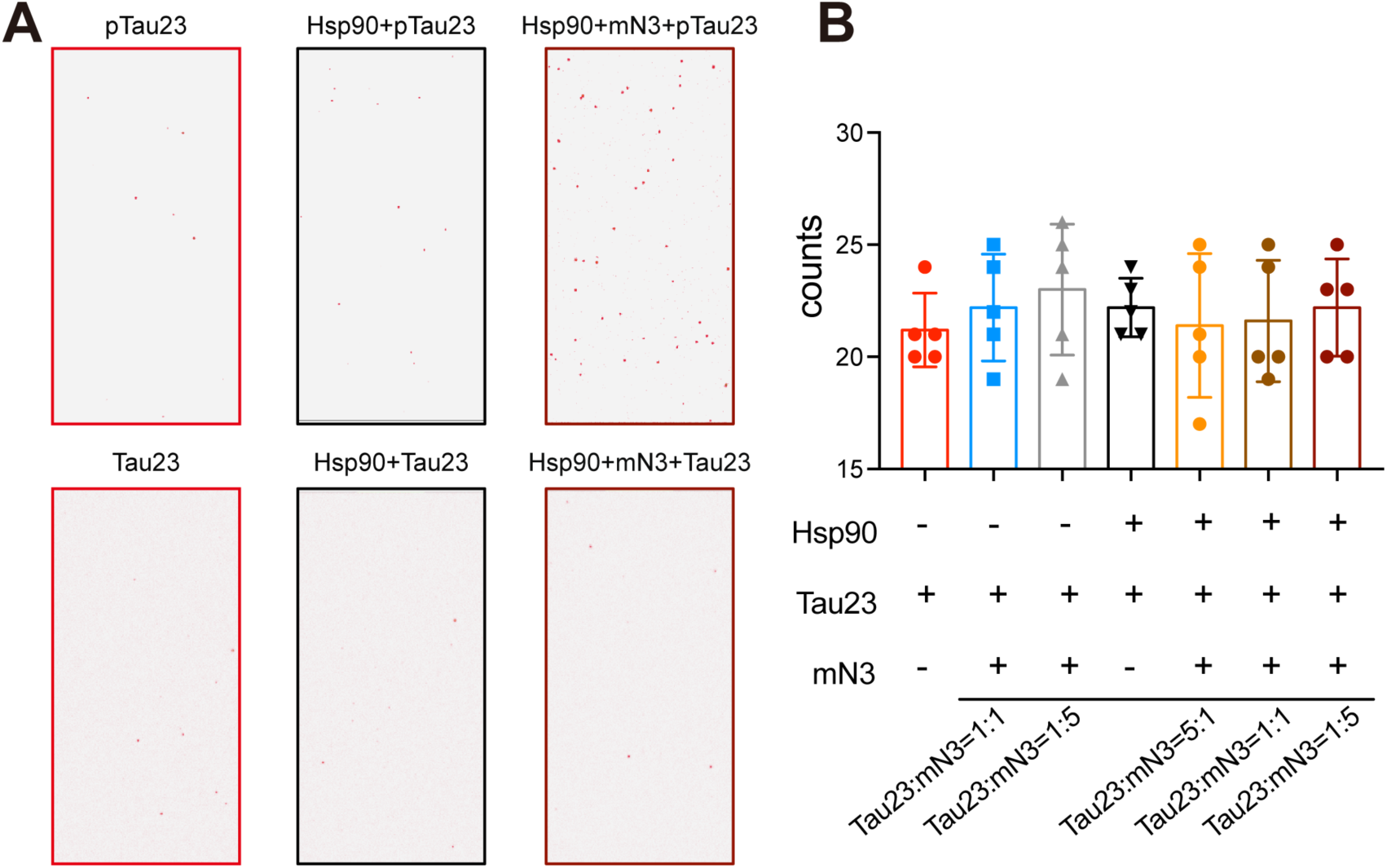
mN3 mediates the binding of pTau23, but not Tau23, to Hsp90. E TIRF microscopic images show the enhancement of mN3 to the binding between pTau23 and Hsp90. F The average number of fluorescent counts per imaging area detected by SMPull. The concentrations of mN3 from left to right are 0, 4, 20, 0, 0.8, 4, and 20 nM. Error bars denote standard deviations (s.d.) (n = 10).

**Supplementary Table 1.**
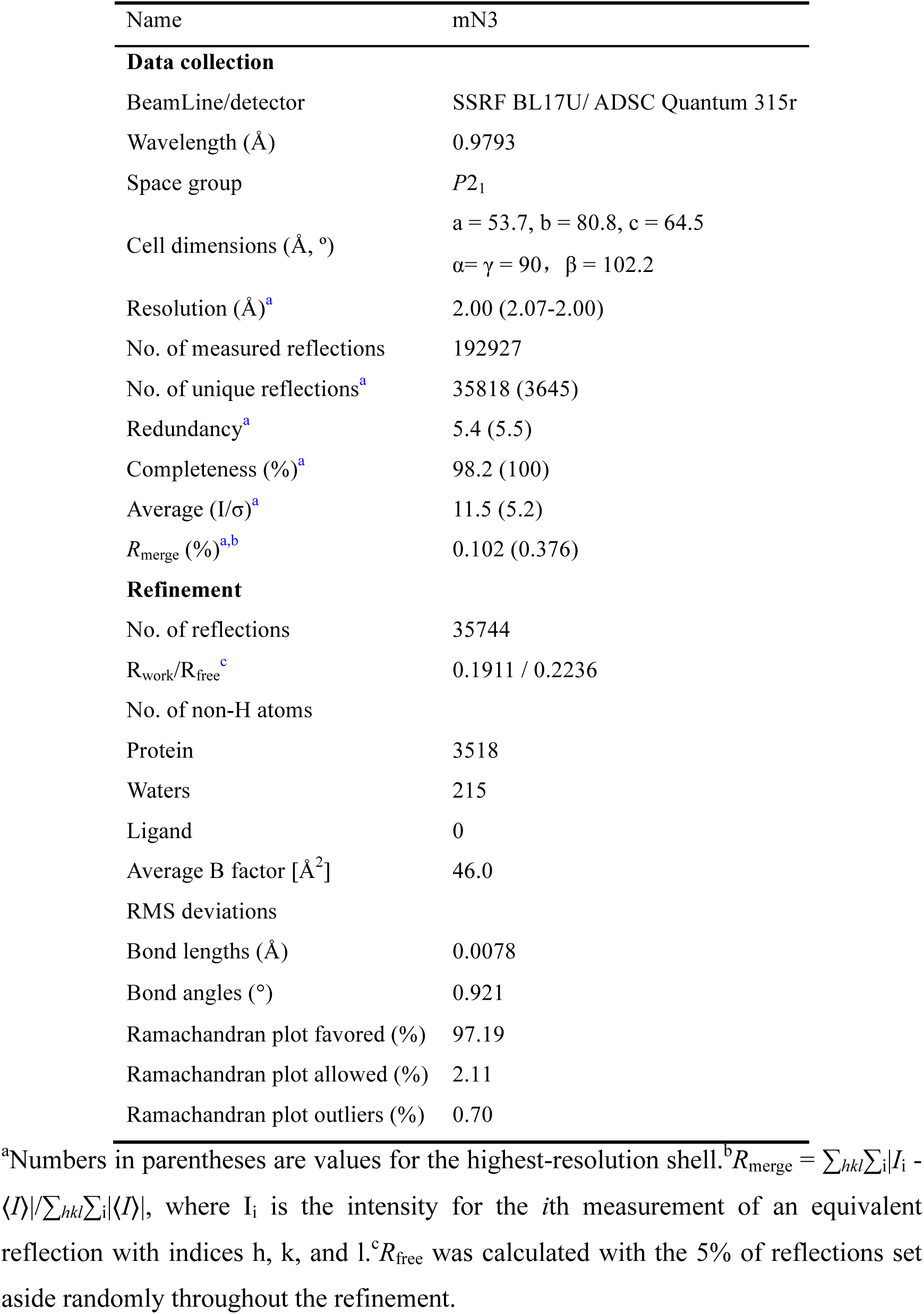
Data collection and structure refinement statistics of mN3.

**Supplementary Table 2.**
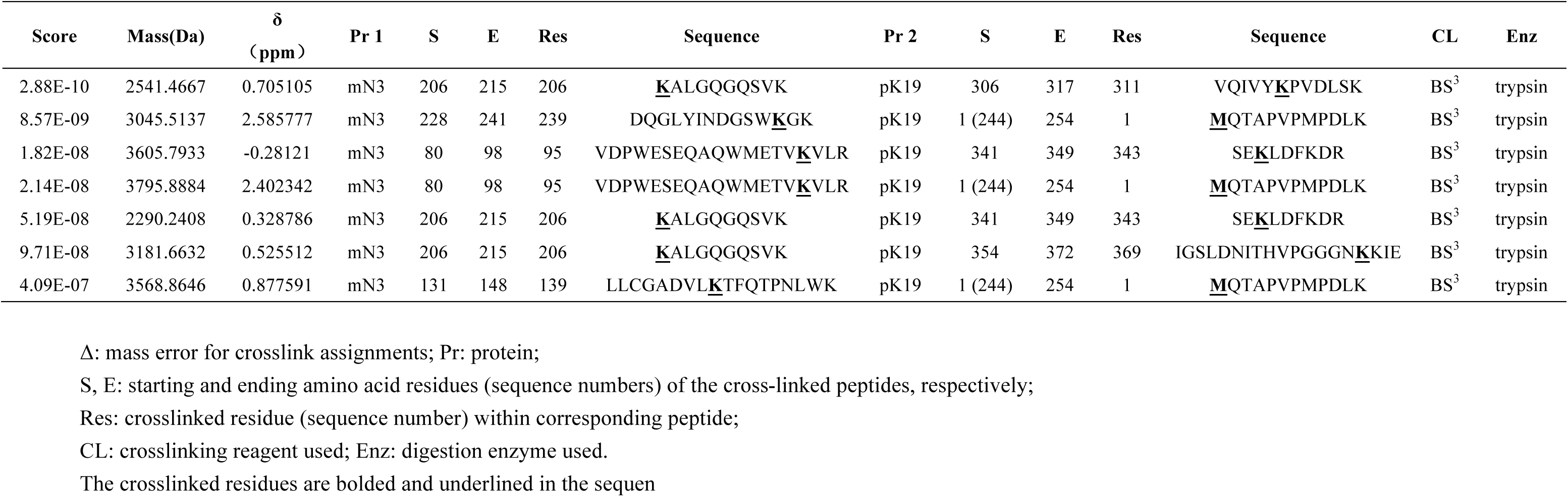
Cross-linked peptides between pK19 and mN3.

